# A gene-regulatory network model for density-dependent and sex-biased dispersal evolution during range expansions

**DOI:** 10.1101/2023.07.18.549508

**Authors:** Jhelam N. Deshpande, Emanuel A. Fronhofer

## Abstract

Dispersal is key to understanding ecological and evolutionary dynamics. Dispersal may itself evolve and exhibit phenotypic plasticity. Specifically, organisms may modulate their dispersal rates in response to the density of their conspecifics (density-dependent dispersal) and their own sex (sex-biased dispersal). While optimal dispersal plastic responses have been derived from first principles, the genetic and molecular basis of dispersal plasticity has not been modelled. An understanding of the genetic architecture of dispersal plasticity is especially relevant for understanding dispersal evolution during rapidly changing spatial ecological conditions such as range expansions. In this context, we develop an individual-based metapopulation model of the evolution of density-dependent and sex-biased dispersal during range expansions. We represent the dispersal trait as a gene-regulatory network (GRN), which can take population density and an individual’s sex as an input and analyse emergent context- and condition-dependent dispersal responses. We compare dispersal evolution and ecological dynamics in this GRN model to a standard reaction norm (RN) approach under equilibrium metapopulation conditions and during range expansions. We find that under equilibrium metapopulation conditions, the GRN model produces emergent density-dependent and sex-biased dispersal plastic response shapes that match the theoretical expectation of the RN model. However, during range expansion, when mutation effects are large enough, the GRN model leads to faster range expansion because GRNs can maintain higher adaptive potential. Our results imply that, in order to understand eco-evolutionary dynamics in contemporary time, the genetic architecture of traits must be taken into account.

## Introduction

Dispersal is key to understanding both ecology and evolution since it impacts the population dynamics of organisms and the distribution of their genes (Ronce, 2007; Govaert et al., 2019). Further, not only may dispersal evolve in response to spatio-temporal variation in fitness expectations, kin structure, and inbreeding avoidance (Bowler and Benton, 2005), but it also exhibits phenotypic plasticity. While it is recognised that dispersal can respond to the internal state (condition-dependent dispersal; Clobert et al. 2009) and the external environment (context-dependent dispersal) of an organism (Fronhofer et al., 2018), the consequences of accounting for underlying molecular and genetic processes that generate dispersal plasticity are unclear (Saastamoinen et al., 2018). In the present study, therefore, we will outline as proof-of-concept how accounting for the genetic basis of dispersal plasticity in models can impact our understanding of dispersal evolution. We focus on two examples of dispersal plastic responses that have been well-studied: density-dependent (Harman et al., 2020) and sex-biased (Li and Kokko, 2019) dispersal.

Dispersal rates of organisms show plastic responses to local population density and may increase (positive density-dependent dispersal), decrease (negative density-dependent dispersal), or even be unimodal (reviewed in Harman et al. (2020)). Theoretical work has focused on the evolution of positive density-dependent dispersal, which evolves when there is negative density-dependence in density regulation (Gyllenberg and Metz, 2001; Poethke and Hovestadt, 2002). If individuals are present in a patch that has a smaller population density than an average patch, they experience less competition and, therefore, tend to stay in their natal patch (no dispersal), and those in patches with higher than average densities tend to leave their natal patch with a probability that increases with local population density due to increased competition (Gyllenberg and Metz, 2001; Poethke and Hovestadt, 2002). Many theoretical studies have assumed different shapes of positive density-dependence: linear (Travis and Dytham, 1999) or sigmoid (Kun and Scheuring, 2006; Bocedi et al., 2012; Travis et al., 2009). However, the theoretical expectation in discrete time models is given by a function in which dispersal is zero below a threshold and then increases in a saturating manner beyond it (Poethke and Hovestadt, 2002). Apart from a first principles justification, this reaction norm shape outcompetes all the others in pairwise competition simulation experiments (Hovestadt et al., 2010). Similarly, sex-biased dispersal is known to evolve due to asymmetry in limiting resources, kin competition, or inbreeding depression (Li and Kokko, 2019). When females mate with a randomly chosen male, this leads to the evolution of male-biased dispersal, that is, males tend to disperse more than females, since they experience greater variability in mates, which is a limiting resource (Gros et al., 2009).

Apart from the first principle approaches already described above (e.g., Poethke and Hovestadt (2002) and Gyllenberg and Metz (2001)), the shape of the optimal dispersal plastic response can also be obtained by other methods. A function value trait approach has been used in which different “loci” represent the trait value corresponding to a given environment (Dieckmann et al., 2006) or differing internal conditions (Gros et al., 2009). Finally, some studies have relied on polynomials if the function-valued trait approach was too computationally demanding (Deshpande et al., 2021). Closer to the present study, Ezoe and Iwasa (1997) standardised a neural network model against analytically derived reaction norms for density-dependent dispersal.

However, fundamentally, these optimal reaction norms must have an underlying molecular and genetic basis (Saastamoinen et al., 2018), that is, there must be a genotype-to-phenotype map (Alberch, 1991; Nichol et al., 2019) that can process internal states and environmental conditions, leading to the emergence of plastic responses at the phenotypic level. One such representation of a genotype-to-phenotype map is the gene-regulatory network (GRN) model proposed by Wagner (1994) and its variants (Spirov and Holloway, 2013). While this is still a highly simplified representation of molecular processes that generate plasticity, gene-regulatory network approaches can reveal how phenotypic plasticity modifies evolvability by introducing developmental constraints (Draghi and Whitlock, 2012; Brun-Usan et al., 2021). For example, under conditions of rapid environmental change, Draghi and Whitlock (2012) modelled phenotypic plasticity of two correlated quantitative traits using a model combining GRN and quantitative genetics approaches. They found that plastic populations, which evolve in heterogeneous environments and have genes that receive an input from the external environment, exhibit evolvability in the direction of environmental variation and adapt most easily. van Gestel and Weissing (2016) modelled bacterial sporulation using a GRN approach, incorporating phenotypic plasticity by allowing the regulatory genes to receive environmental inputs, and found that a GRN approach allows for greater diversity in the response to novel conditions than a classical reaction norm approach, capturing a greater adaptive potential.

Thus, one context in which accounting for molecular mechanisms for dispersal plasticity may be relevant is understanding rapid evolution during directional change, such as during range expansions (Miller et al., 2020). How quickly organisms spread in space depends, besides reproduction, centrally on dispersal. Since dispersal has a genetic basis (Saastamoinen et al., 2018) and can evolve (Ronce, 2007), the potentially rapid evolution of dispersal ability can impact range expansion dynamics, but, vice versa, range expansions can also drive dispersal evolution by spatial sorting and selection, wherein more dispersive individuals end up at the range expansion front (Shine et al., 2011). It has also been shown that the speed of range expansions depends critically on whether dispersal increases or decreases with population density (Altwegg et al., 2013). In theoretical work, density-dependent dispersal can lead to accelerating range expansions (Travis et al., 2009), due to the evolution of decreased positive density-dependence of dispersal at range fronts. Yet, experimental studies have shown both, reductions (Fronhofer et al., 2017; Dahirel et al., 2021, 2022) and increases (Mishra et al., 2020) in positive density-dependence of dispersal during range expansion.

Building on this work, we posit that gene-regulatory networks can be used to model dispersal plasticity. Using this bottom-up approach, we here seek to understand whether processes at the molecular level, particularly gene regulation, yield a similar plastic response to the theoretically predicted optimal reaction norm (Poethke and Hovestadt, 2002) in the case of density-dependent dispersal. Hence, we develop an individual-based metapopulation model, in which dispersal can evolve to be plastic to local population density. We represent the genetic architecture of density-dependent dispersal using a GRN that takes as an input the local population density, regulatory genes process this input and finally output a continuous dispersal probability trait. We compare the GRN model to the theoretically expected reaction norm (RN) shape proposed by Poethke and Hovestadt (2002). Finally, we also investigate whether such a match to theoretical expectations holds if dispersal can additionally be sex-biased (Li and Kokko, 2019). To highlight how the genetic architecture of dispersal plasticity impacts predictions under conditions of rapid change, we model range expansions.

Thus, in this study, we address the following questions: 1) Does a more mechanistic GRN model of plasticity lead to the emergence of what is predicted from first principles at the RN level? 2) What are the ecological and evolutionary consequences of a more complex but mechanistic model under native equilibrium metapopulation conditions and during range expansions?

### Model description

#### General description

We develop a discrete-time and discrete-space individual-based metapopulation model of a sexually reproducing diploid species in which dispersal can evolve and be plastic to local population density and sex. Density regulation is local within a patch of the metapopulation, and local dynamics follow a Beverton-Holt model of logistic growth (Beverton and Holt, 1957). We represent the genetic basis of an individual’s dispersal trait by a Wagner-like (Wagner, 1994) gene-regulatory network (GRN), that takes as input and processes population density as an external cue and sex as an internal state, producing as an output its dispersal probability (Fig. 1 A, C). In order to compare our model to the theoretically expected plastic response in the cases of density-dependent dispersal and density-dependent and sex-biased dispersal, we develop additional models (Fig. 1 B, D) using the reaction norm approach described in Poethke and Hovestadt (2002).

**Figure 1:**
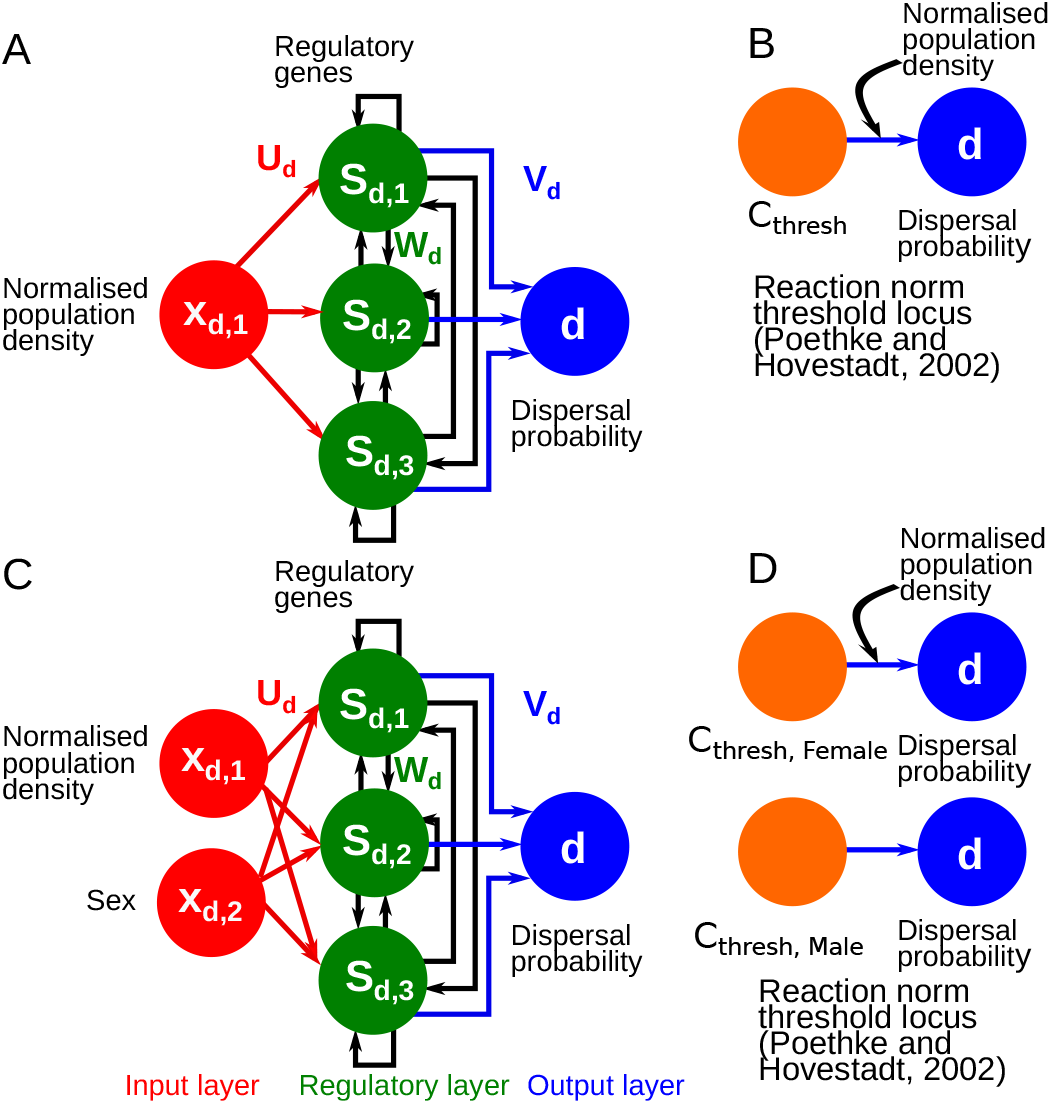
GRN (A, C) and RN (B, D) models for density-dependent (A–B) and density-dependent and sex-biased dispersal (C–D). The assumed GRN model has an input layer, which is a vector **x**_*d*_ of external states or external cues, in our case, population density alone (A) and population density and sex (B). The regulatory genes receive this input via the input weights ***U*** _*d*_. Genes have expression states denoted by **S**_*d*_, and interactions between these genes are encoded by a regulatory matrix ***W*** _*d*_. The effects of these genes are encoded by the matrix ***V*** _*d*_. In the case of density-dependent dispersal, the RN model is represented by a single quantitative locus, which is the threshold of the function derived by Poethke and Hovestadt (2002), and for density-dependent and sex biased dispersal, two loci with sex dependent expression encode the threshold. We compare the evolution of dispersal plasticity and range expansion dynamics between the reaction norm and GRN approaches.

Individuals are initially present in the central 10 × 5 patches of out of a 500 × 5 grid landscape, for 20000 generations (time-steps), in order for the dispersal genotypes to reach (quasi)-equilibrium. We assume that these individuals can start range expansion in the *x*-dimension after 20000 generations. Therefore, the boundary conditions in the *x*-direction are reflecting for the first 20000 generations. In the y-direction, boundary conditions are toroidal, hence the landscape resembles a hollow tube. Range expansions can take place till the expanding population has moved 245 patches from the central 10 × 5 patches in either direction along the *x*-dimension. Range expansions stop when the expanding population reaches the boundary of the landscape in the *x*-dimension.

### Life cycle

#### Dispersal

We assume that dispersal is natal. The probability that an individual disperses is given by its genetically encoded plastic response to local population density (Fig. 1 A–B) alone or local population density and sex (Fig. 1 C–D). The plastic response may either be encoded by a GRN or the threshold (Fig. 1 A, C) of a theoretically expected reaction norm (Fig. 1 B, D). If an individual disperses, one of the eight nearest neighbouring patches (Moore neighbourhood) is chosen as the target patch. Dispersal costs (Bonte et al., 2012) are captured by the dispersal mortality *µ*, which is the probability that an individual dies while dispersing.

#### Reproduction and inheritance

After dispersal, individuals reproduce sexually. The population dynamics in a patch follow the Beverton-Holt model of logistic growth (Beverton and Holt, 1957):

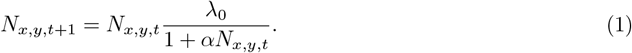

Here, *λ*_0_ is the intrinsic growth rate, and *α* is the intra-specific competition coefficient. This model reaches an expected equilibrium density of 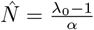 in the absence of spatial structure for *λ*_0_ > 1. A female first chooses a mate at random, and then produces a number of offspring drawn from a Poisson distribution with a mean 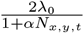. The factor of 2 corrects for the fact that only females reproduce and keeps *λ*_0_ interpretable at the population level. The offspring inherit the alleles to the various parameters of the GRN, or the threshold of the theoretically expected reaction norm, one from each parent at each locus. In the GRN model, we assume that the per locus per allele mutation rate decreases linearly from *m*_*max*_ = 0.1 to *m*_*min*_ = 0.0001 in the first 5000 time steps and is constant after (Deshpande and Fronhofer, 2022). Since the GRN model has a large number of parameters, using larger mutation rates initially allows the fitness landscape to be coarsely explored quickly without the trait value getting stuck in a local optimum. In the RN model, *m*_*min*_ = *m*_*max*_ = 0.0001 throughout the simulation. The mutation effects per allele per locus for both models are drawn from Gaussian distribution with a standard deviation *σ*_*m*_ = 0.1.

Generations are non-overlapping, therefore, the offspring generation replaces the parental generation. In addition, we assume that there may be random patch extinctions every generation with a probability of *ϵ* per patch. These extinctions represent density-independent, catastrophic external impacts.

### Gene-regulatory network (GRN) model

The genetic basis of density-dependent and sex-biased dispersal is modelled by a modified Wagner-like (Wagner, 1994) gene-regulatory network model (Deshpande and Fronhofer, 2022). We assume that the dispersal probability *d* results from the linear combination (Draghi and Whitlock, 2012) of equilibrium gene expression states 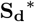 of *n* = 4 genes that interact with each other. We assume that organisms can detect local population density (Fellous et al., 2012; Fronhofer et al., 2015) and their own sex, which can produce a plastic response in their gene expression, hence, their dispersal trait. Thus, these genes take as input the population density normalised by the expected equilibrium density of the Beverton-Holt model 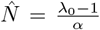 and sex (0 and 1 for female and male, respectively) of an organism (Fig. 1 A, C). The gene-regulatory network has three layers: an input layer (**x**_*d*_; taking population density and sex as cues), a regulatory layer (**S**_**d**_(*I*); vector of gene expression states corresponding to an iteration *I* of the developmental process), and an output layer (*d*; the dispersal probability trait) (van Gestel and Weissing, 2016). These layers are connected to each other by matrices of weights: the input weights (***U*** _*d*_), regulatory weights (***W*** _*d*_) and output weights (***V*** _*d*_). The expression state of a gene is a sigmoid function of the input it receives from the environment and other genes (Siegal and Bergman, 2002) and can take values between −1 and 1. Each gene has its own properties: a slope (**r**_*d*_) and a threshold (***θ***_*d*_) to this sigmoid. The slopes and thresholds of all genes, along with the elements of the input, regulatory, and output weight matrices, are encoded by a diploid locus each with two alleles. The mid parental value at each locus is used to iterate through gene expression states according to equation Eq. 2.

Thus, the developmental process for the dispersal trait is characterised by the following difference equation (Deshpande and Fronhofer, 2022) where **S**_**d**_(*I*) is the vector of gene expression states for *n* genes and *m* inputs at each iteration of the developmental process:

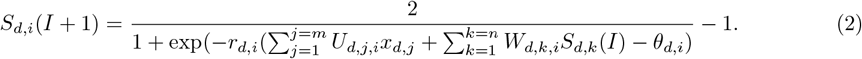

The equilibrium gene expression states 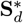 are obtained after *I* = 20 iterations. Individuals with GRNs that do not reach steady state equilibrium at this point die (Wagner, 1994). The dispersal probability is then calculated as the linear combination of these equilibrium gene expression states (Draghi and Whitlock, 2012) as:

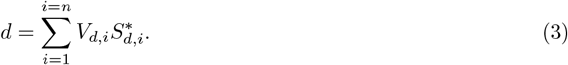

### Reaction norm (RN) model

We compare the plastic response that arises in the GRN model to the theoretically expected optimal reaction norm (RN) derived from first principles for density-dependent dispersal (Poethke and Hovestadt, 2002). In discrete time metapopulation models with logistic growth, dispersal probability is expected to be 0 below a threshold local population density and increase in a saturating manner with it. Thus, dispersal probability *d* is given by:

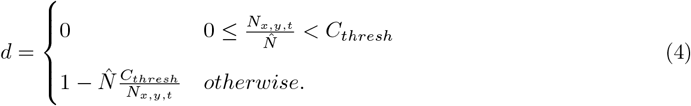

Here, 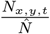 is the local population density normalised by the expected equilibrium population density, and *C*_*thresh*_ is the threshold density, which can be optimised by simulations (Poethke and Hovestadt, 2002). Thus, in the RN model, we assume that the threshold density *C*_*thresh*_ is genetically encoded by a single diploid locus with two alleles. Individuals detect local population density *N*_*x,y,t*_ and disperse with a probability given by equation Eq. 4.

We also extend this approach to sex-biased and density-dependent dispersal by encoding two different threshold normalised densities as two loci, *C*_*thresh,M*_ and *C*_*thresh,F*_. *C*_*thresh,M*_ is expressed if the individual is male, and *C*_*thresh,F*_ is expressed if the individual is female.

### Analysis

We analyse both GRN and RN models (Fig. 1) for density-dependent dispersal (GRN DDD and RN DDD) alone and for density-dependent and sex-biased dispersal (GRN DDD + sex bias and RN DDD + sex bias). Model parameters are found in Table 1. Since dispersal evolution ultimately is driven by costs and benefits, we run 50 replicate model simulations for dispersal mortality *µ* ∈ {0.01, 0.1, 0.3} and a random patch extinction risk of *ϵ* ∈ {0, 0.05, 0.1}. We first compare the long term (*t* = 20000 time steps) evolved plastic response in the GRN DDD and GRN DDD + sex bias models to the expected optimal reaction norms RN DDD and RN DDD + sex bias models under standard metapopulation conditions. After 20000 time steps, individuals begin range expansions, and we compare range expansion speeds between the GRN and RN models. In order to test the sensitivity of our results to assumed mutation rates we run additional simulations for the GRN DDD model with mutation effects that are 1*/*4 times the standard GRN DDD model (termed the GRN DDD small mutation effects model).

**Table 1:** Model Parameters/Variables.

| Model Parameter/Variable | Description | Values |
| --- | --- | --- |
| $N_{x,y,t}$ | Population density in the patch $x, y$ at time $t$ | dynamical |
| $\lambda_0$ | Intrinsic growth rate in Beverton-Holt model | 2 |
| $\alpha$ | Intra-specific competition coefficient in Beverton-Holt model | 0.01 |
| $\mu$ | Dispersal mortality | 0.01, 0.1, 0.3 |
| $\epsilon$ | Local patch extinction probability | 0, 0.05, 0.1 |
| $m_{min}$ | Mutation rate at equilibrium | 0.0001 |
| $m_{max}$ | Mutation rate at the beginning | 0.1 for GRN, $m_{max} = m_{min}$ for others |
| $\sigma_m$ | Effect size (standard deviation) of mutations | 0.1 |
| $\mathbf{x}$ | Vector of inputs to the GRN | in simulation |
| $n$ | Number of regulatory genes | 4 |
| $\mathbf{S}_{d,I}$ | Vector of expression states of regulatory genes at iteration I | in simulation |
| $\mathbf{U}_d$ | $m \times n$ matrix with each element $U_{ji}$ representing the connection between the input $j$ and gene $i$ | evolves, initialised from a normal distribution with $sd = 1$ |
| $\mathbf{W}_d$ | $n \times n$ matrix with each element $W_{ki}$ representing the connection between the gene $k$ and gene $i$ | evolves, initialised from a normal distribution with $sd = 1$ |
| $\mathbf{V}_d$ | $1 \times n$ matrix with each element $V_i$ representing the connection between the gene $k$ and the output | evolves, initialised from a normal distribution with $sd = 1$ |
| $\boldsymbol{\theta}_d$ | Thresholds of regulatory genes | evolves, initialised from a normal distribution with $sd = 1$ |
| $\mathbf{r}_d$ | Slopes of regulatory genes | evolves, initialised from a normal distribution with $sd = 1$ |
| $C_{thresh}$ | Threshold for density-dependent dispersal in RN model | evolves, initialised from a uniform distribution between 0 to 1 |
| $C_{thresh,F}$ | Threshold for density-dependent and sex-biased dispersal in RN model for females | evolves, initialised from a uniform distribution between 0 to 1 |
| $C_{thresh,M}$ | Threshold for density-dependent and sex-biased dispersal in RN model for males | evolves, initialised from a uniform distribution between 0 to 1 |

## Results and discussion

### Evolution of the density-dependent dispersal plastic response in the GRN and RN models

The density-dependent dispersal plastic response (Fig. 2) obtained after 20000 generations in the GRN DDD model matches the theoretically expected optimum (RN DDD; Poethke and Hovestadt (2002)) most closely for high extinction probability (for *ϵ* = 0.05 and 0.1) and high dispersal mortality (for *µ* = 0.1 and 0.3). When there are no patch extinctions (for *ϵ* = 0), the GRN DDD plastic response differs from the theoretical optimum likely because the individuals in the metapopulation are not exposed to a wide range of population densities, preventing optimisation (Fig. 2). Finally, low dispersal mortality (*µ* = 0.01) also reduces optimisation. This is likely because the strength of selection for reduced dispersal is low since the fitness cost of a non-optimal dispersal decision is low. In addition to Fig. 2, the quality of optimisation in the GRN DDD model is assessed in SI Fig. S1, which also shows that the GRN DDD model is closest to the theoretical optimum under conditions of high patch extinctions and dispersal mortality. Our result that optimisation in the GRN DDD model is least effective under conditions of low dispersal mortality and extinction probability is consistent with those of Hovestadt et al. (2010) who show that other strategies can co-exist with the theoretically expected optimal response (Poethke and Hovestadt, 2002) in competition experiments under similar conditions of low environmental variability and low dispersal mortality.

**Figure 2:**
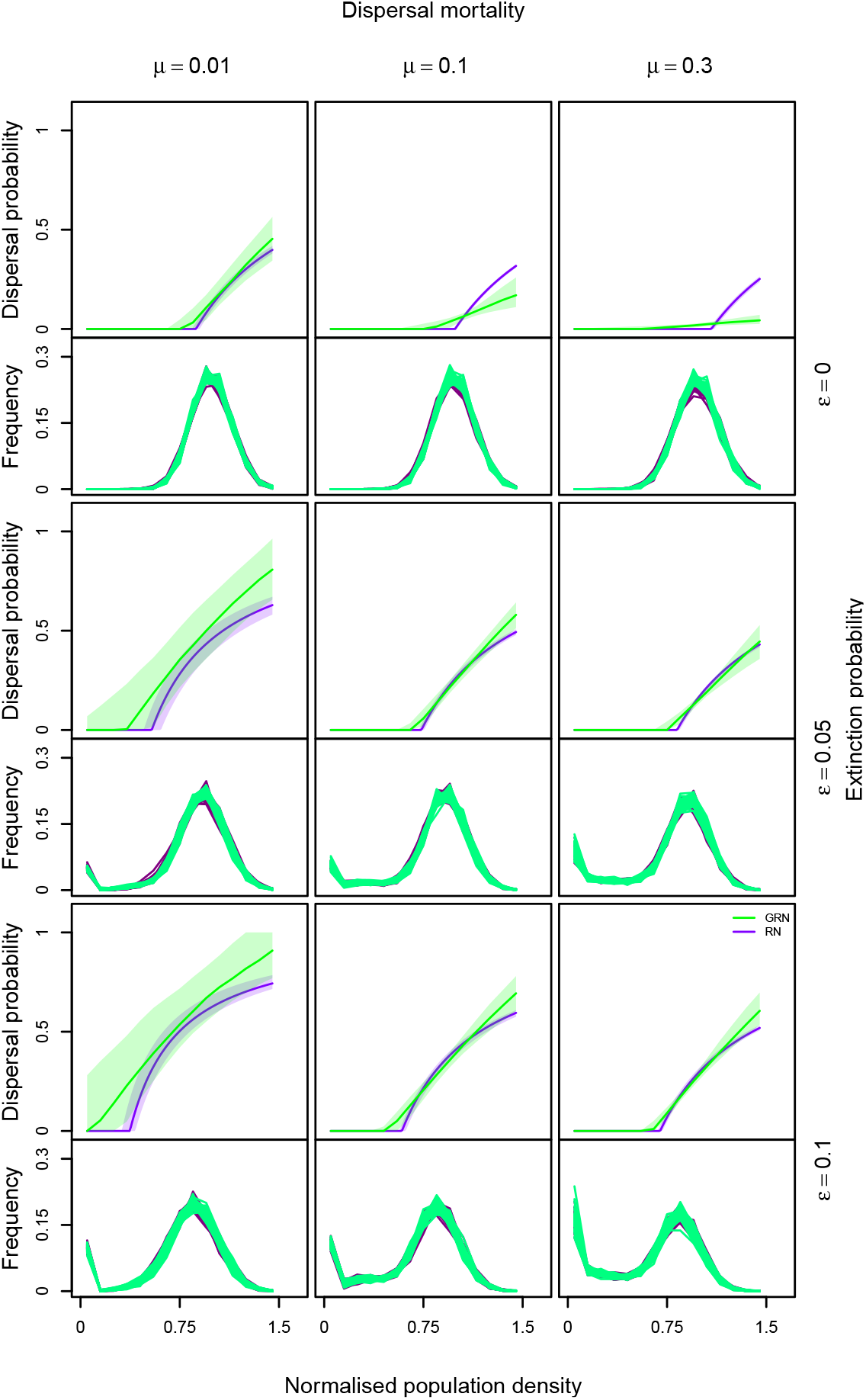
The GRN DDD (green) matches the RN DDD (purple) model for high extinction probability and dispersal mortality. The match is greatest at population densities that are most frequent. Dispersal mortality increases from left to right (*µ* ∈ {0.01, 0.1, 0.3}), from top to bottom, extinction probability increases (*ϵ*∈ { 0, 0.05, 0.1 }). For each combination of dispersal mortality and extinction probability, the evolutionarily stable (ES) dispersal probability and the histogram of population densities that occur during the simulation are plotted for both GRN and RN models. ES dispersal probability as a function of population density normalised by the expected equilibrium population density (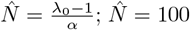 in the present study). The purple line represents the density-dependent dispersal plastic response calculated from the median threshold *C*_*thresh*_ obtained after 20000 time steps over all individuals, and the shaded region from the inter-quartile range in the RN DDD model. The green line represents the calculated median GRN output for 1000 randomly chosen individuals pooled across all 50 replicates at the end of 20000 time steps. Fixed parameters: intrinsic growth rate: *λ*_0_ = 2, intraspecific competition coefficient: *α* = 0.01, and number of regulatory genes: *n* = 4.

The amount and direction of phenotypic variation that is maintained in the gene-regulatory network model, again depends on dispersal mortality and extinction probability. Particularly, this variation is comparable in the GRN DDD and the RN DDD models at high dispersal mortality and extinction probability, but at low dispersal mortality, greater phenotypic variation is maintained in the GRN DDD model (SI Fig. S2). This is because of the evolution of greater sensitivity to mutation relative to the RN DDD model (SI Fig. S4) when dispersal mortality is low, which is expected since the negative fitness consequences of a non-optimal dispersal decision increase with increasing dispersal mortality. Reduced optimisation (SI Fig. S1) and increased phenotypic variation (SI Fig. S2) in the GRN DDD model under conditions of low dispersal mortality and extinction probability do not seem to have important consequences on metapopulation dynamics since the distribution of observed population densities in both models do not differ (Fig. 2). To test whether this maintenance of variation at low population densities is a consequence of assumed mutation rates we also simulated a GRN DDD model with a quarter of the mutation effects (0.25*σ*_*m*_; GRN DDD smaller effects model). We find that phenotypic variation maintained at low population densities is still higher in the GRN DDD smaller effects model relative to the RN DDD model but quantitatively lower than the GRN DDD model (see Fig. S3). The ecoevolutionary consequences of the differences in maintenance of phenotypic variation will be explored in detail below.

In summary, the GRN DDD model produces a plastic response similar to theoretical expectation (Poethke and Hovestadt, 2002) when the strength of selection on dispersal is sufficiently high and the individuals across generations are exposed to a wide range of population densities, that is, when dispersal mortality and extinction probability are high. Deviations from this expectation occur when the strength of selection on dispersal is low (at low dispersal mortality and extinction probability) and when individuals across generations are not exposed to a wide range of population densities.

### Evolution of the density-dependent and sex-biased dispersal plastic response in the GRN and RN models

Dispersal may not only depend on the external context but also on internal conditions (Clobert et al., 2009), such as the sex of the potentially dispersing individual. Fig. 3 shows that including the input of an internal condition, sex, along with local population density, as explored in the previous subsection, leads to the emergence of a density-dependent and sex-biased dispersal plastic response in the GRN DDD + sex bias model similar to the optimal response in the RN DDD + sex bias model. The conditions of dispersal mortality and extinction probability for greater optimisation of the GRN model with sex-bias are similar to those when dispersal is not sex-biased (SI Fig. S5). Similar to the scenario in which dispersal is not sex-biased, greater phenotypic variation (SI Fig. S6) and sensitivity to mutation (SI Fig. S7) occur at low dispersal mortality. These differences in phenotypic variation in dispersal reaction norms do not have consequences on the distribution of population density in the metapopulation (SI Fig. S9)

**Figure 3:**
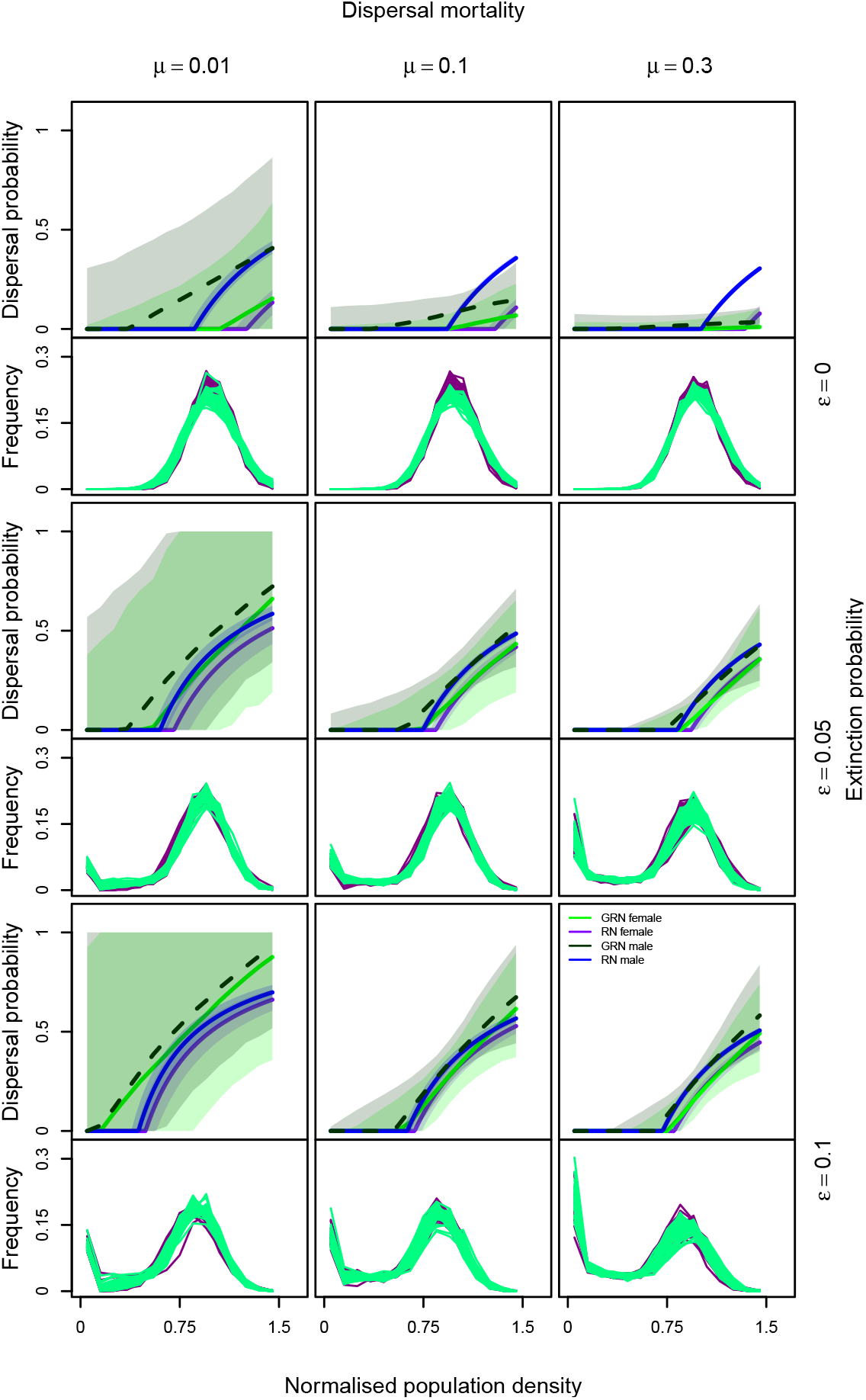
The density-dependent dispersal and sex-biased plastic response in the GRN DDD + sex bias model matches the RN DDD + sex bias model at high dispersal mortality and extinction probability. The match is greatest at population densities that are most frequent. Dispersal mortality increases from left to right (*µ* ∈ {0.01, 0.1, 0.3}), from top to bottom, extinction probability increases (*ϵ* ∈ {0, 0.05, 0.1}). For each combination of dispersal mortality and extinction probability, the evolutionarily stable (ES) dispersal probability and the histogram of population densities that occur during the simulation are plotted for both GRN and RN models. ES dispersal probability as a function of population density normalised by the expected equilibrium population density (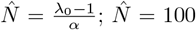 in the present study). The blue and purple lines represent the ES reaction norms for males and females in the RN model from Poethke and Hovestadt (2002). The dark green and green lines represent the calculated GRN output for 1000 randomly chosen individuals at the end of 20000 time steps corresponding to male and female sex respectively. The transparency of the points is weighted by the frequency of occurrence of population density so as to only represent the GRN plastic response for those densities that occur during the simulation. Fixed parameters: *λ*_0_ = 2 and *α* = 0.01. Number of regulatory genes *n* = 4.

Focusing on sex-bias, the density-dependent dispersal threshold is lower for males than for females, leading to male-biased dispersal in our simulations. This is consistent with previous work on sex-biased dispersal, which shows that males experience greater stochasticity in mate finding, which leads to the evolution of greater dispersal in males relative to females (Gros et al., 2009).

### Genetic architecture of dispersal plasticity impacts eco-evolutionary dynamics of range expansion

Under equilibrium metapopulation conditions, we have shown that both density-dependent dispersal and sex-biased dispersal plastic responses readily evolve in gene-regulatory network models and outline the conditions in which they match their theoretical optimum. But what are the ecological consequences of such plastic responses under novel conditions? In order to answer this question, after 20000 time steps, we allow for range expansions in both the GRN and RN models. We find that range expansion speeds are greater in the GRN model overall when local density alone (Fig. 4) and both local density and sex, define dispersal decisions (Fig. 5). In general, the difference between range expansion dynamics in the two models is greater when dispersal mortality is low and the rate of external patch extinctions is high (Fig. 4–5).

**Figure 4:**
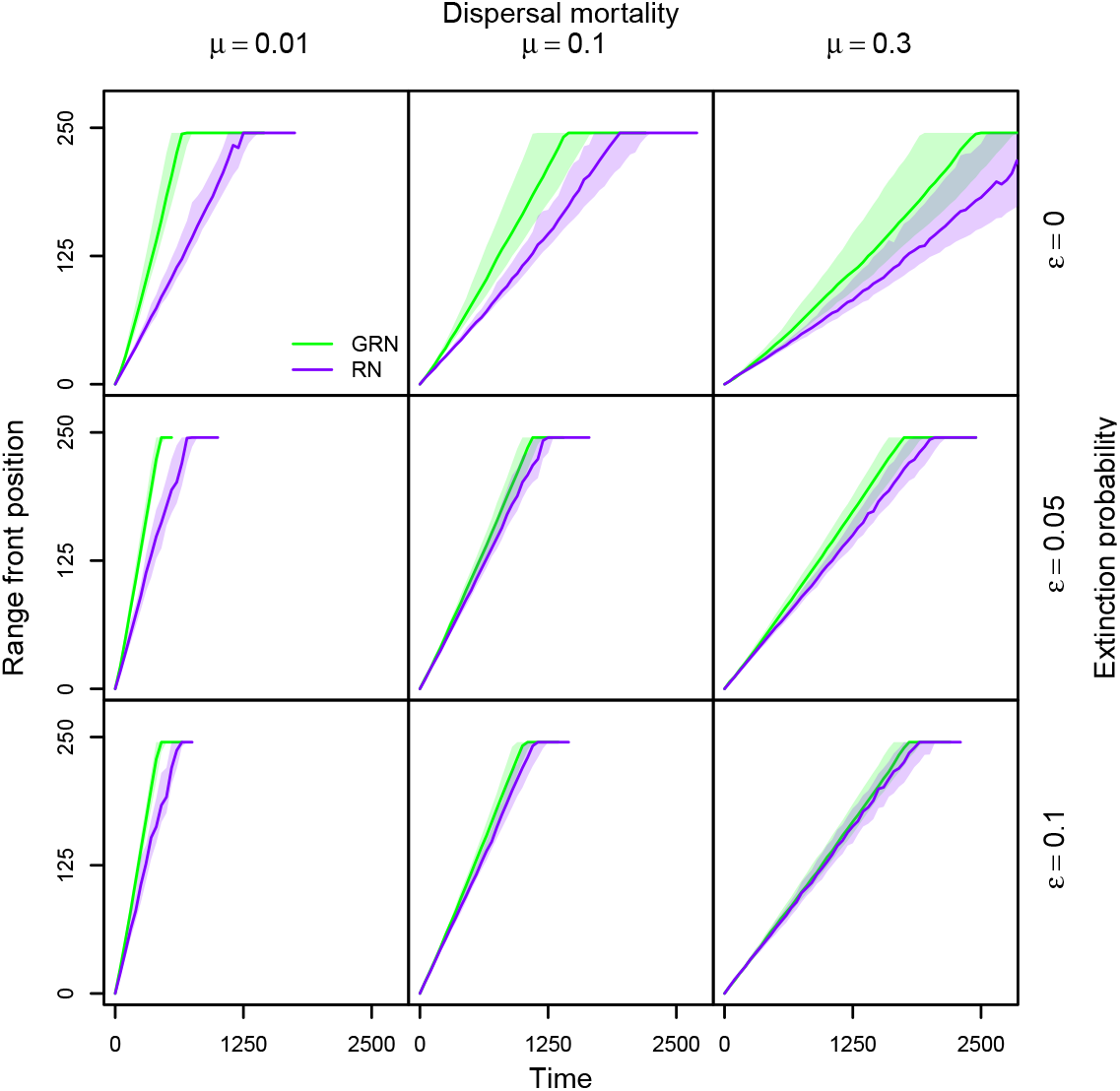
Range expansion dynamics in GRN vs. RN model for DDD. Dispersal mortality increases from left to right (*µ* ∈ { 0.01, 0.1, 0.3}), from top to bottom, extinction probability increases (*ϵ* ∈ { 0, 0.05, 0.1}). We plot the median and quartiles of range front position as a function of time for the GRN model and RN model. The range front is defined as the farthest occupied patch from the range core. Fixed parameters: *λ*_0_ = 2 and *α* = 0.01. Number of regulatory genes: *n* = 4.

**Figure 5:**
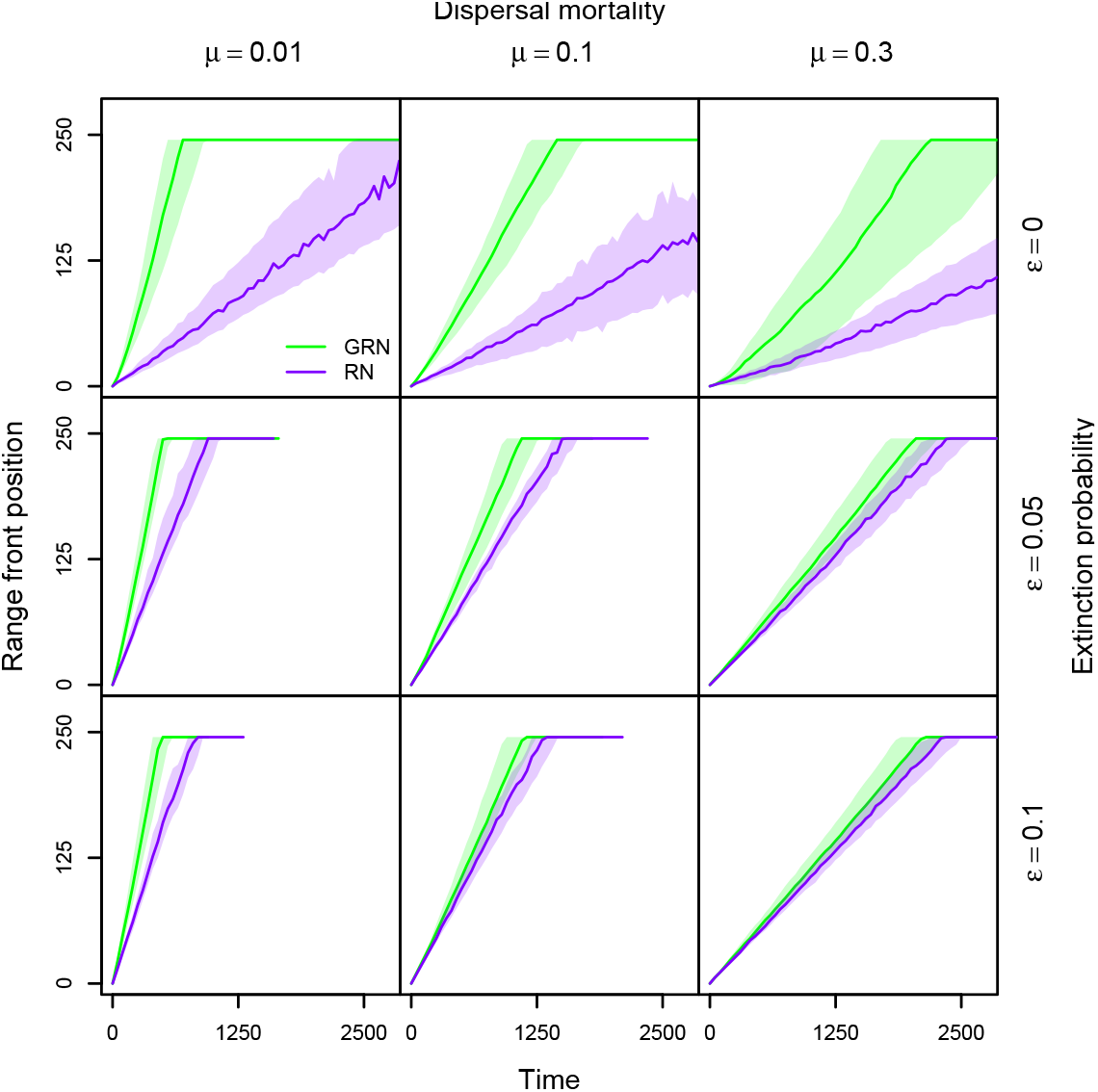
Range expansion dynamics in GRN vs. RN model for DDD and sex bias. Dispersal mortality increases from left to right (*µ* ∈ { 0.01, 0.1, 0.3 }), from top to bottom, extinction probability increases (*ϵ* ∈ { 0, 0.05, 0.1}). We plot the median and quartiles of range front position as a function of time for the GRN model and RN model. The range front is defined as the farthest occupied patch from the range core. Fixed parameters: *λ*_0_ = 2 and *α* = 0.01. Number of regulatory genes: *n* = 4.

These patterns of faster range expansion speeds in the GRN model and the conditions of low dispersal mortality and extinction probability that produce them can be understood on the basis of the evolutionary history of the metapopulation before range expansions begin. As seen in the previous section, the GRN model maintains greater phenotypic variation (SI Fig. S2 and S6) under conditions of low dispersal mortality (see SI Fig. S8–S9 for individual reaction norms). Moreover, when there are no patch extinctions, variation is also maintained at low population densities since these population densities do not occur during the equilibrium metapopulation phase, allowing for the accumulation of genetic variation (Fig. S8–S9). This variation is then spatially sorted (Shine et al., 2011), leading to the evolution of greater dispersal rates at the range expansion front in the GRN model relative to the RN model (SI Fig. S10–S11). This is evident in the trajectories of evolved dispersal as a function of time in the range front (SI Fig. S12) in which the GRN model leads to the evolution of greater dispersal rates early on in the range expansion indicating that standing genetic variation is being sorted. Travis et al. (2009) have previously shown that accelerating invasions can be found in models assuming sigmoid density-dependent dispersal reaction norms. They argue that this allows them to have a relatively flexible function, where not just a threshold, as in Poethke and Hovestadt (2002), but also other properties of the reaction norm can evolve. We reconcile the two approaches because, at the equilibrium metapopulation level without assuming a particular shape of the plastic response, on average, the shape that emerges is the one predicted by Poethke and Hovestadt (2002) but the GRN approach has greater evolutionary flexibility as in Travis et al. (2009).

Our results that the GRN model leads to faster range expansions are sensitive to the assumption of smaller mutation effects. In simulations where the per locus per allele mutation effects are smaller (= 0.25*σ*_*m*_; GRN DDD small mutation effects), while the amount of phenotypic variation maintained at low population density is quantitatively lower than the RN DDD model and higher than the GRN DDD model, this does not translate to faster range expansions (see SI Fig. S13). This indicates that the per locus per allele mutation effects in the GRN model need to be high enough for a minimal amount of phenotypic variation to be maintained at low population densities. Interestingly, the possibility of sexbiased and density-dependent dispersal increases the difference between the dynamics of the RN model and the GRN model. Generally, male-biased dispersal (Fig. 3) slows down range expansions (Miller et al., 2011) due to the fact that males cannot reproduce by themselves, implying that population, hence range expansion dynamics, are female-limited. Thus, the availability of variation at densities that do not occur in equilibrium metapopulation conditions in the GRN model further amplifies differences between the two models relative to density-dependent dispersal alone.

### General discussion

In summary, we developed a model for density-dependent and sex-biased dispersal that assumes that dispersal results from the effects of a gene-regulatory network. We find that under conditions that are experienced in equilibrium metapopulations, the emergent predicted plastic response matches existing theoretical predictions well for conditions of high dispersal mortality and extinction probability. We then compare range expansion dynamics between a GRN and an RN model and find that the GRN model leads to faster range expansions if mutation effects are large because of the maintenance of greater variation when selection on dispersal is not high and the population has relatively stable dynamics (no patch extinctions).

The theoretical literature usually uses highly simplified representations of the genetic architecture of traits like dispersal, most often only representing them at the level of the phenotype (Saastamoinen et al., 2018). Particularly, adaptive dynamics approaches (Parvinen et al., 2006), which assume small mutation effects and rare mutations, allow for optimal traits or reaction norms to be derived, analytically or by means of simulation, as a function of ecological equilibria (Govaert et al., 2019). Quantitative genetics approaches may further highlight constraints on optimisation of reaction norms such as genetic correlations (Gomulkiewicz and Kirkpatrick, 1992). Further, in simulations similar to ours, one quantitative locus with additive effects is often assumed (Saastamoinen et al., 2018). On the other hand, studies of genetic architecture rarely make ecological conditions explicit, with an abstract representation of selection on traits by assuming a fitness function that is *a priori* defined rather than a result of underlying ecological processes (e.g., studies using the Wagner model; Wagner 1994). Few studies highlight the advantage of incorporating both explicit ecological dynamics and genetic architectures. A notable exception is, for example, van Gestel and Weissing (2016), who compare GRN and RN approaches for bacterial sporulation and show the GRN approaches maintain greater diversity of plastic responses which makes them more evolvable under novel conditions.

In our study, we recapture the theoretically expected and known phenotypic relationships between population density and dispersal (Poethke and Hovestadt, 2002), confirming the validity of our approach. Importantly, under novel, low-density conditions experienced during range expansions, the differences observed between expansion dynamics in the different models make clear that approaches based on reaction norms may not be able to predict eco-evolutionary dynamics under novel conditions.

Our results underline the relevance of understanding genetic architecture (Yamamichi, 2022) for ecoevolutionary dynamics (Melián et al., 2018; Fronhofer et al., 2023), particularly for dispersal (Saastamoinen et al., 2018) and its response to internal and external cues (Clobert et al., 2009). While empirical evidence supporting our work is scarce, Brisson et al. (2010) showed differences in gene expression between winged and un-winged phenotypes of pea aphids, particularly in their wing development gene-regulatory network. In this system, winged morphs are often induced due to crowding, and the relative production of dispersive and non-dispersive (reproductive) females depends on developmental cues, including crowding. More generally, our GRN approach can be used to understand how dispersal responds to other internal (e.g., infection state; Iritani and Iwasa 2014 or body condition; Baines et al. 2020) and external cues, for example, the presence of parasites (Deshpande et al., 2021) or predators (Poethke et al., 2010).

Our study links very closely to Ezoe and Iwasa (1997), who used a neural network to compare the evolution of dispersal reaction norms to analytical predictions. They showed that the neural network was able to produce plastic responses similar to the analytically derived reaction norm while finding some consistent deviations from this optimal response. In our study, we go beyond these results by highlighting the conditions of dispersal mortality and extinction probability that yield reaction norms closest to the expected optimal response. Moreover, using a gene-regulatory network approach allows us to place our work in context of previous work investigating the relationship between phenotypic plasticity and evolvability (Draghi and Whitlock, 2012; van Gestel and Weissing, 2016; Brun-Usan et al., 2021).

GRN models and models of GP maps often use highly abstract representations of the environment (for example, Draghi and Whitlock 2012) and gene expression as the phenotype directly under selection (for example, Espinosa-Soto et al. 2011). These approaches have been useful in defining, for example, how evolvability of phenotypes is linked with phenotypic plasticity (van Gestel and Weissing, 2016) and the alignment between genetic, environmental perturbations, and direction of selection, and how this impacts evolvability in multi-trait systems (Draghi and Whitlock, 2012; Brun-Usan et al., 2021).

However, in an eco-evolutionary framework (Govaert et al., 2019; Fronhofer et al., 2023), ecological interactions define selection on a trait. Ecological dynamics also define the trait that is under selection. Therefore, considering gene expression as a phenotype directly under selection may not always be appropriate, and gene expression state to phenotype maps must be included (Chevin et al., 2022). This is relevant because, for example, the association of extremes of gene expression (Rünneburger and Rouzic, 2016) with increased mutational sensitivity (decreased robustness) is actually reversed (Deshpande and Fronhofer, 2022). Further, while such a map is likely to be more complex than our assumed linear gene expression to phenotype map, approaches such as ours and that of van Gestel and Weissing (2016) also narrow the range of possible environments under native conditions and also help define phenotypes under selection that are ecologically informed.

The latter point becomes clear when considering our results on range expansion dynamics. Taking into account both genetic architecture and the ecological conditions that shape the evolution of dispersal plasticity, the GRN model leads to the maintenance of variation in conditions (densities) that are not very frequent under equilibrium metapopulation conditions. This variation is then spatially sorted (Shine et al., 2011) during range expansion. However, in the RN approach, this maintenance of variation under equilibrium metapopulation conditions does not happen since only the threshold to the reaction norm is under selection. We see the consequences of the spatial sorting of dispersal in the fact that range expansions are generally faster in GRN approaches, when dispersal is density-dependent alone, and sex bias only increases the difference between the two models. This has previously been discussed in the literature as a form of cryptic variation, particularly “hidden reaction norms” (Schlichting, 2008), which represent differences in genotypes that are not normally expressed at the phenotypic level but might be expressed if the genotype is perturbed due to mutation or recombination, but also when the environment is perturbed. Our results are similar to the findings of van Gestel and Weissing (2016) who showed that in their GRN model, the release of cryptic variation in native environments can lead to more adaptive plastic responses in novel conditions.

We additionally show that these mechanisms driving differences in range expansion dynamics may critically depend on assumed mutation effects. This is because a comparison between the GRN and RN model is not straightforward since the sensitivity of the dispersal reaction norm to mutations is determined both by the underlying genetic architecture (e.g., GRN vs. RN) and the assumed per locus per allele mutation effects at the loci encoding the plastic response. While under conditions in which there is greater selection on dispersal and relatively heterogeneous conditions of population density (high dispersal mortality and extinction probability), similar behaviour between GRN and RN model is predicted, at conditions in which selection on dispersal is not as strong (low dispersal probability and extinction probability) sufficient variation may not be maintained to speed up range dynamics relative to an RN model.

More importantly, the GRN model also provides a molecular-mechanistic basis for plasticity. While the GRN is likely to be more complicated in reality, the different layers of the gene-regulatory network that produce the plastic response can be interpreted biologically. For example, the input layer represents the external environmental cue, population density, which can be sensed as, for example, the reduced availability of resources or other chemical and mechanical cues (Fellous et al., 2012; Fronhofer et al., 2015) resulting from a larger local density of individuals. The regulatory layer can be interpreted as the gene expression states in cells of a relevant developmental stage that respond to local population density. Empirical studies of gene regulation in a dispersal context remain rare. Yagound et al. (2022) have shown gene expression differences using mRNA sequencing in the brains of the invasive Australian cane toad in a few genes. In their study, dispersal-related genes generally showed elevated expression at the range front. In this system, associated life history and physiological changes are particularly well studied in terms of range expansion dynamics (Phillips et al., 2006; Perkins et al., 2013). Other examples include wing polyphenism in pea aphids (Brisson et al., 2010), and dispersal in yellow-bellied marmots (Armenta et al., 2019). This relative scarcity of empirical studies, together with the relatively important effects predicted by our model, clearly call for more work, both empirical and theoretical, to understand how genotype-to-phenotype maps impact eco-evolutionary dynamics.

## Author contributions

JND and EAF conceived the study. JND developed and analysed the models in collaboration with EAF. JND wrote the manuscript in collaboration with EAF.

## Acknowledgements

This is publication ISEM-YYYY-XXX of the Institut des Sciences de l’Evolution – Montpellier.

## Data availability

Simulation code is available via GitHub and Zenodo (DOI: https://doi.org/10.5281/zenodo.8160132).

## Notes

https://doi.org/10.5281/zenodo.8160132

### Competing Interest Statement

The authors have declared no competing interest.

### Summary of Updates

The previous version of the manuscript was not the final revised version recommended by PCI. The current one is the correct one with the badge.

### Funding

- Agence Nationale de la Recherche: ANR-19-CE02-0015

## References

Alberch, P. (1991). From genes to phenotype: dynamical systems and evolvability. Genetica, 84(1):5–11.

Altwegg, R., Collingham, Y. C., Erni, B., and Huntley, B. (2013). Density-dependent dispersal and the speed of range expansions. Divers. Distrib., 19(1):60–68.

Armenta, T. C., Cole, S. W., Geschwind, D. H., Blumstein, D. T., and Wayne, R. K. (2019). Gene expression shifts in yellow-bellied marmots prior to natal dispersal. Behav. Ecol., 30(2):267–277.

Baines, C. B., Travis, J. M. J., McCauley, S. J., and Bocedi, G. (2020). Negative density-dependent dispersal emerges from the joint evolution of density- and body condition-dependent dispersal strategies. Evolution, 74(10):2238–2249.

Beverton, R. J. H. and Holt, S. J. (1957). On the dynamics of exploited fish populations. Chapman & Hall, London.

Bocedi, G., Heinonen, J., and Travis, J. M. J. (2012). Uncertainty and the role of information acquisition in the evolution of context-dependent emigration. Am. Nat., 179(5):606–620.

Bonte, D., Van Dyck, H., Bullock, J. M., Coulon, A., Delgado, M., Gibbs, M., Lehouck, V., Matthysen, E., Mustin, K., Saastamoinen, M., Schtickzelle, N., Stevens, V. M., Vandewoestijne, S., Baguette, M., Barton, K., Benton, T. G., Chaput-Bardy, A., Clobert, J., Dytham, C., Hovestadt, T., Meier, C. M., Palmer, S. C. F., Turlure, C., and Travis, J. M. J. (2012). Costs of dispersal. Biol. Rev., 87(2):290–312.

Bowler, D. E. and Benton, T. G. (2005). Causes and consequences of animal dispersal strategies: relating individual behaviour to spatial dynamics. Biol. Rev., 80(2):205–225.

Brisson, J. A., Ishikawa, A., and Miura, T. (2010). Wing development genes of the pea aphid and differential gene expression between winged and unwinged morphs. Insect Mol. Biol., 19:63–73.

Brun-Usan, M., Rago, A., Thies, C., Uller, T., and Watson, R. A. (2021). Development and selective grain make plasticity ‘take the lead’ in adaptive evolution. BMC Ecol. Evol., 21(1):205.

Chevin, L.-M., Leung, C., Le Rouzic, A., and Uller, T. (2022). Using phenotypic plasticity to understand the structure and evolution of the genotype–phenotype map. Genetica, 150(3):209–221.

Clobert, J., Galliard, J.-F. L., Cote, J., Meylan, S., and Massot, M. (2009). Informed dispersal, heterogeneity in animal dispersal syndromes and the dynamics of spatially structured populations. Ecol. Lett., 12(3):197–209.

Dahirel, M., Bertin, A., Calcagno, V., Duraj, C., Fellous, S., Groussier, G., Lombaert, E., Mailleret, L., Marchand, A., and Vercken, E. (2021). Landscape connectivity alters the evolution of density-dependent dispersal during pushed range expansions. bioRxiv.

Dahirel, M., Guicharnaud, C., and Vercken, E. (2022). Individual variation in dispersal, and its sources, shape the fate of pushed vs. pulled range expansions. bioRxiv.

Deshpande, J. N. and Fronhofer, E. A. (2022). Genetic architecture of dispersal and local adaptation drives accelerating range expansions. Proc. Natl. Acad. Sci. U. S. A., 119(31):e2121858119.

Deshpande, J. N., Kaltz, O., and Fronhofer, E. A. (2021). Host–parasite dynamics set the ecological theatre for the evolution of state- and context-dependent dispersal in hosts. Oikos, 130(1):121–132.

Dieckmann, U., Heino, M., and Parvinen, K. (2006). The adaptive dynamics of function-valued traits. J. Theor. Biol., 241(2):370–389.

Draghi, J. A. and Whitlock, M. C. (2012). Phenotypic plasticity facilitates mutational variance, genetic variance, and evolvability along the major axis of environmental variation. Evolution, 66(9):2891–2902.

Espinosa-Soto, C., Martin, O. C., and Wagner, A. (2011). Phenotypic plasticity can facilitate adaptive evolution in gene regulatory circuits. BMC Evol. Biol., 11(1).

Ezoe, H. and Iwasa, Y. (1997). Evolution of condition-dependent dispersal: A genetic-algorithm search for the ESS reaction norm. Population Ecology, 39(2):127–137.

Fellous, S., Duncan, A., Coulon, A., and Kaltz, O. (2012). Quorum Sensing and Density-Dependent Dispersal in an Aquatic Model System. PLOS One, 7(11):e48436.

Fronhofer, E. A., Corenblit, D.; Deshpande, J. N., Govaert, L., Huneman, P., Viard, F., Jarne, P., and Puijalon, S. (2023). Eco-evolution from deep time to contemporary dynamics: the role of timescales and rate modulators. Ecol. Lett.

Fronhofer, E. A., Gut, S., and Altermatt, F. (2017). Evolution of density-dependent movement during experimental range expansions. J. Evol. Biol., 30(12):2165–2176.

Fronhofer, E. A., Kropf, T., and Altermatt, F. (2015). Density-dependent movement and the consequences of the Allee effect in the model organism Tetrahymena. J. Anim. Ecol., 84(3):712–722.

Fronhofer, E. A., Legrand, D., Altermatt, F., Ansart, A., Blanchet, S., Bonte, D., Chaine, A., Dahirel, M., De Laender, F., De Raedt, J., et al. (2018). Bottom-up and top-down control of dispersal across major organismal groups. Nat. Ecol. Evol., 2(12):1859.

Gomulkiewicz, R. and Kirkpatrick, M. (1992). Quantitative genetics and the evolution of reaction norms. Evolution, 46(2):390.

Govaert, L., Fronhofer, E. A., Lion, S., Eizaguirre, C., Bonte, D., Egas, M., Hendry, A. P., De Brito Martins, A., Melián, C. J., Raeymaekers, J. A. M., Ratikainen, I. I., Saether, B.-E., Schweitzer, J. A., and Matthews, B. (2019). Eco-evolutionary feedbacks—Theoretical models and perspectives. Funct. Ecol., 33(1):13–30.

Gros, A., Poethke, H. J., and Hovestadt, T. (2009). Sex-specific spatio-temporal variability in reproductive success promotes the evolution of sex-biased dispersal. Theor. Popul. Biol., 76:13–18.

Gyllenberg, M. and Metz, J. A. J. (2001). On fitness in structured metapopulations. J. Math. Biol., 43(6):545–560.

Harman, R. R., Goddard, J., Shivaji, R., and Cronin, J. T. (2020). Frequency of occurrence and population-dynamic consequences of different forms of density-dependent emigration. Am. Nat., 195(5):851–867.

Hovestadt, T., Kubisch, A., and Poethke, H.-J. (2010). Information processing in models for density-dependent emigration: A comparison. Ecol. Model., 221(3):405–410.

Iritani, R. and Iwasa, Y. (2014). Parasite infection drives the evolution of state-dependent dispersal of the host. Theor. Popul. Biol., 92:1–13.

Kun, A. and Scheuring, I. (2006). The evolution of density-dependent dispersal in a noisy spatial population model. Oikos, 115(2):308–320.

Li, X.-Y. and Kokko, H. (2019). Sex-biased dispersal: a review of the theory. Biol. Rev., 94(2):721–736.

Melián, C. J., Matthews, B., de Andreazzi, C. S., Rodríguez, J. P., Harmon, L. J., and Fortuna, M. A. (2018). Deciphering the interdependence between ecological and evolutionary networks. Trends Ecol. Evol., 33(7):504–512.

Miller, T. E., Angert, A. L., Brown, C. D., Lee-Yaw, J. A., Lewis, M., Lutscher, F., Marculis, N. G., Melbourne, B. A., Shaw, A. K., Szucs, M., et al. (2020). Eco-evolutionary dynamics of range expansion. Ecology, 101(10):e03139.

Miller, T. E. X., Shaw, A. K., Inouye, B. D., and Neubert, M. G. (2011). Sex-biased dispersal and the speed of two-sex invasions. Am. Nat., 177(5):549–561.

Mishra, A., Chakraborty, P. P., and Dey, S. (2020). Dispersal evolution diminishes the negative density dependence in dispersal. Evolution, 74(9):2149–2157.

Nichol, D., Robertson-Tessi, M., Anderson, A. R. A., and Jeavons, P. (2019). Model genotype–phenotype mappings and the algorithmic structure of evolution. J. R. Soc. Interface, 16(160):20190332.

Parvinen, K., Dieckmann, U., and Heino, M. (2006). Function-valued adaptive dynamics and the calculus of variations. Journal of Mathematical Biology, 52(1):1–26.

Perkins, T. A., Phillips, B. L., Baskett, M. L., and Hastings, A. (2013). Evolution of dispersal and life history interact to drive accelerating spread of an invasive species. Ecol. Lett., 16(8):1079–1087.

Phillips, B. L., Brown, G. P., Webb, J. K., and Shine, R. (2006). Invasion and the evolution of speed in toads. Nature, 439(7078):803–803.

Poethke, H., Weisser, W., and Hovestadt, T. (2010). Predator-Induced Dispersal and the Evolution of Conditional Dispersal in Correlated Environments. Am. Nat., 175(5):577–586.

Poethke, H. J. and Hovestadt, T. (2002). Evolution of density–and patch–size–dependent dispersal rates. Proc. R. Soc. B-Biol. Sci., 269(1491):637–645.

Ronce, O. (2007). How does it feel to be like a rolling stone? ten questions about dispersal evolution. Annu. Rev. Ecol. Evol. Syst., 38(1):231–253.

Rünneburger, E. and Rouzic, A. L. (2016). Why and how genetic canalization evolves in gene regulatory networks. BMC Evol. Biol., 16(1).

Saastamoinen, M., Bocedi, G., Cote, J., Legrand, D., Guillaume, F., Wheat, C. W., Fronhofer, E. A., Garcia, C., Henry, R., Husby, A., et al. (2018). Genetics of dispersal. Biol. Rev., 93(1):574–599.

Schlichting, C. D. (2008). Hidden reaction norms, cryptic genetic variation, and evolvability. Ann N Y Acad Sci, 1133(1):187–203.

Shine, R., Brown, G. P., and Phillips, B. L. (2011). An evolutionary process that assembles phenotypes through space rather than through time. Proc. Natl. Acad. Sci. U. S. A., 108(14):5708–5711.

Siegal, M. L. and Bergman, A. (2002). Waddington’s canalization revisited: Developmental stability and evolution. Proc. Natl. Acad. Sci. U. S. A., 99(16):10528–10532.

Spirov, A. and Holloway, D. (2013). Using evolutionary computations to understand the design and evolution of gene and cell regulatory networks. Methods, 62(1):39–55.

Travis, J. M. J. and Dytham, C. (1999). Habitat persistence, habitat availability and the evolution of dispersal. Proc R Soc Lond B Biol Sci, 266(1420):723–728.

Travis, J. M. J., Mustin, K., Benton, T. G., and Dytham, C. (2009). Accelerating invasion rates result from the evolution of density-dependent dispersal. J. Theor. Biol., 259(1):151–158.

van Gestel, J. and Weissing, F. J. (2016). Regulatory mechanisms link phenotypic plasticity to evolvability. Sci. Rep., 6(1):24524.

Wagner, A. (1994). Evolution of gene networks by gene duplications: a mathematical model and its implications on genome organization. Proc. Natl. Acad. Sci. U. S. A., 91(10):4387–4391.

Yagound, B., West, A. J., Richardson, M. F., Selechnik, D., Shine, R., and Rollins, L. A. (2022). Brain transcriptome analysis reveals gene expression differences associated with dispersal behaviour between range-front and range-core populations of invasive cane toads in australia. Mol. Ecol., 31(6):1700–1715.

Yamamichi, M. (2022). How does genetic architecture affect eco-evolutionary dynamics? A theoretical perspective. Philos. Trans. R. Soc. B, 377(1855):20200504.

