## Supplementary information for "A gene-regulatory network model for density-dependent and sex-biased dispersal evolution during range expansions"

### Supplementary figures

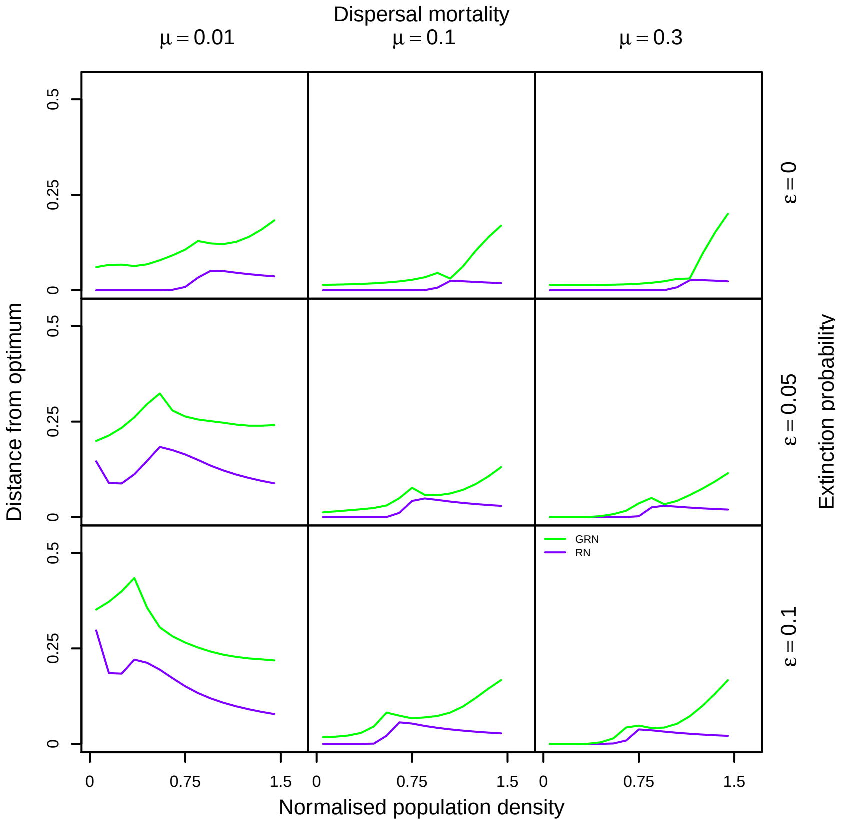


Figure S1: Average distance from optimal plastic response as function of normalised population density for GRN and RN models for DDD. Dipersal mortality increases from left to right ($\mu \in \{0.01, 0.1, 0.3\}$), from top to bottom, extinction probability increases ($\epsilon \in \{0, 0.05, 0.1\}$). The optimal plastic response is calculated from the median of the evolved threshold $C_{thresh}$ in the RN model. The plastic response for 1000 randomly chosen individuals in the GRN and RN models at end of the equilibrium metapopulation phase (20000 time steps) are evaluated at different normalised population densities $0, 0.1, ...1.5$, and the root mean squared distance is calulculated as a measure of deviation from this optimum. We find that overall the deviation from the optimal plastic response is greater in the GRN model relative to the RN model. As dispersal mortality and extinction probability increase, the deviation from optimal plastic response decreases in the GRN model and converges to the RN model. Fixed parameters: $\lambda_0 = 2$ and $\alpha = 0.01$. Number of regulatory genes $n = 4$.

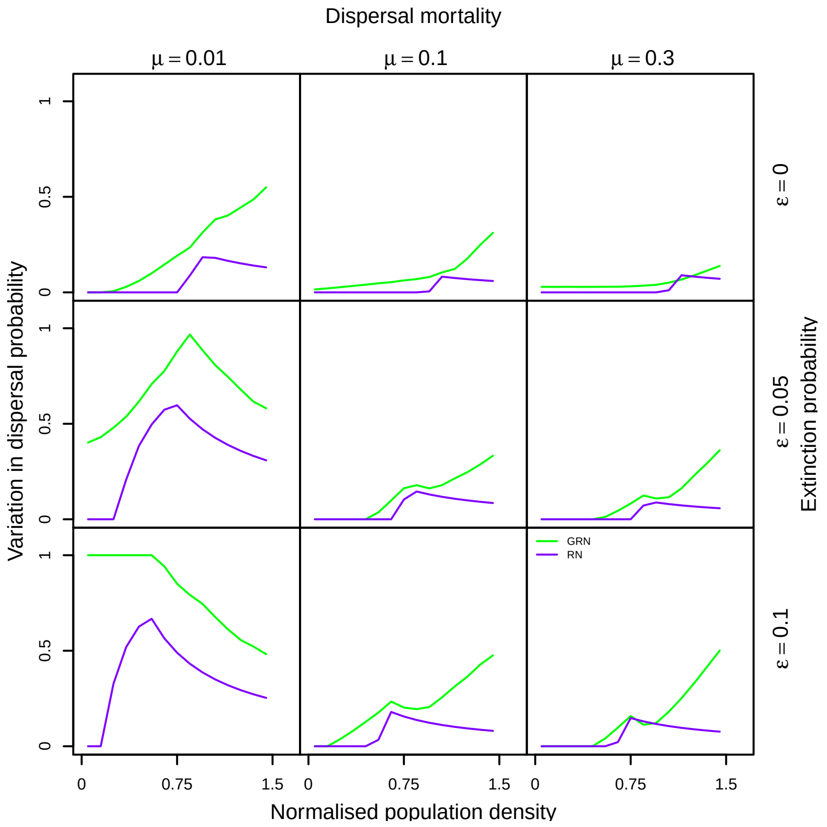


Figure S2: Phenotypic variation maintained in the GRN vs. RN model for DDD as a function of normalised population density. Dipersal mortality increases from left to right ($\mu \in \{0.01, 0.1, 0.3\}$), from top to bottom, extinction probability increases ($\epsilon \in \{0, 0.05, 0.1\}$). Difference between the 95th and 5th percentile in dispersal phenotype as a function of normalised population density plotted for the GRN and RN model. Fixed parameters: $\lambda_0 = 2$ and $\alpha = 0.01$. Number of regulatory genes $n = 4$.

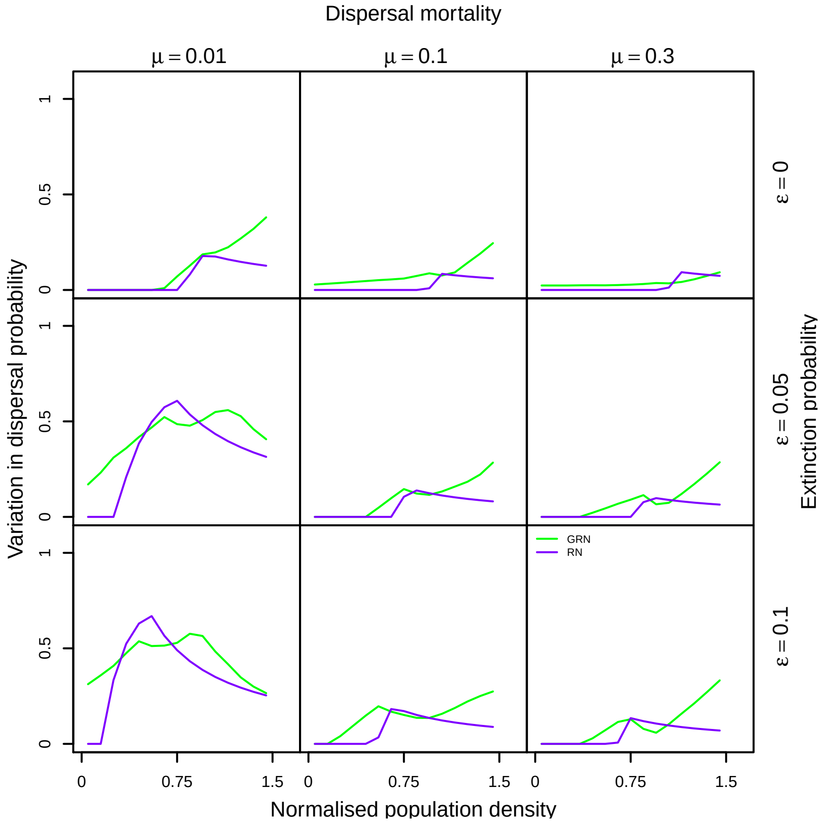


Figure S3: Consequences of lower per locus per allele mutation effects ($0.25\sigma_m$) on phenotypic variation maintained in the GRN vs. RN model for DDD as a function of normalised population density. Dipersal mortality increases from left to right ($\mu \in \{0.01, 0.1, 0.3\}$), from top to bottom, extinction probability increases ($\epsilon \in \{0, 0.05, 0.1\}$). Difference between the 95th and 5th percentile in dispersal phenotype as a function of normalised population density plotted for the GRN and RN model. Phenotypic variation in the GRN DDD model is greater at low population densities relative to the RN model even when mutation effects are small. Fixed parameters: $\lambda_0 = 2$ and $\alpha = 0.01$. Number of regulatory genes $n = 4$.

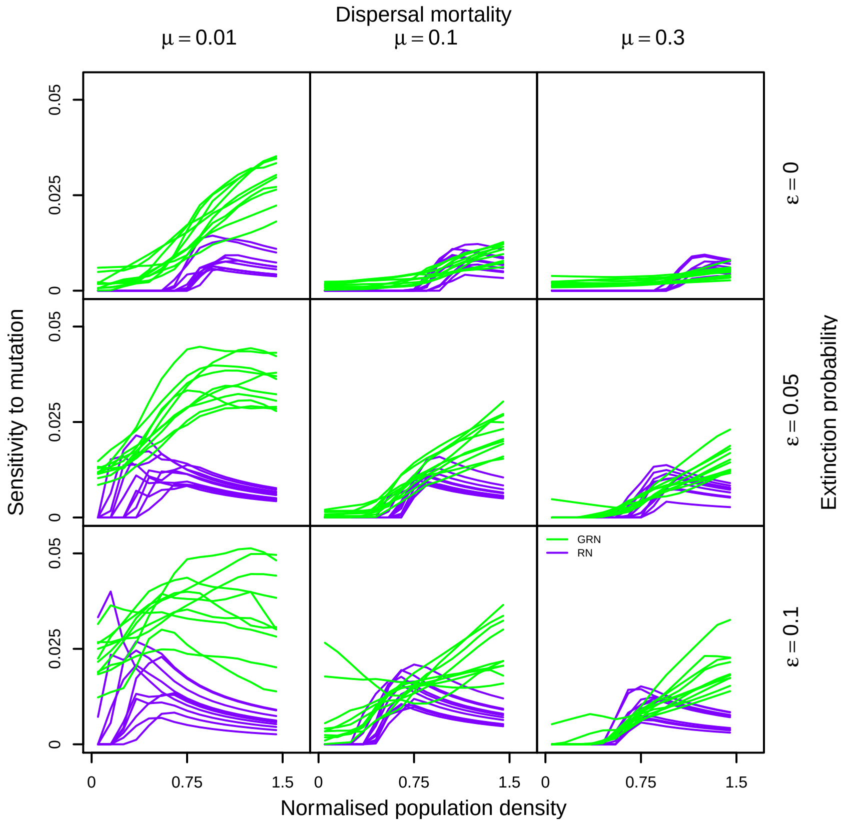


Figure S4: Sensitivity to mutation in the GRN and RN model for DDD as a function of normalised population density. In both models, 1000 individual genotypes are sampled from the last time step of the equilibrium metapopulation phase ($t = 20000$). Dipersal mortality increases from left to right ($\mu \in \{0.01, 0.1, 0.3\}$), from top to bottom, extinction probability increases ($\epsilon \in \{0, 0.05, 0.1\}$). In the RN model, a perturbation drawn from a Gaussian distribution with mean 0 and standard deviation 0.1 is added to the evolved threshold $C_{thresh}$ with probability 0.01. In the GRN model, a perturbation with the same mean and standard deviation is added to to an individual's locus with probability 0.01. This makes both models comparable, since per locus perturbation rate and effect are the same. The sensitivity to mutation is then calculated as the root mean squared difference between the phenotype evaluated from the perturbed and unperturbed genotype. The green and purple lines represent the sensitivity to mutation corresponding to a given normalised population density for 10 replicates of the sampling procedure described above. We find that in the GRN model, sensitivity to mutation is greater at low dispersal mortality and becomes comparable to the RN model as dispersal mortality and extinction probability increases. Fixed parameters: $\lambda_0 = 2$ and $\alpha = 0.01$. Number of regulatory genes $n = 4$.

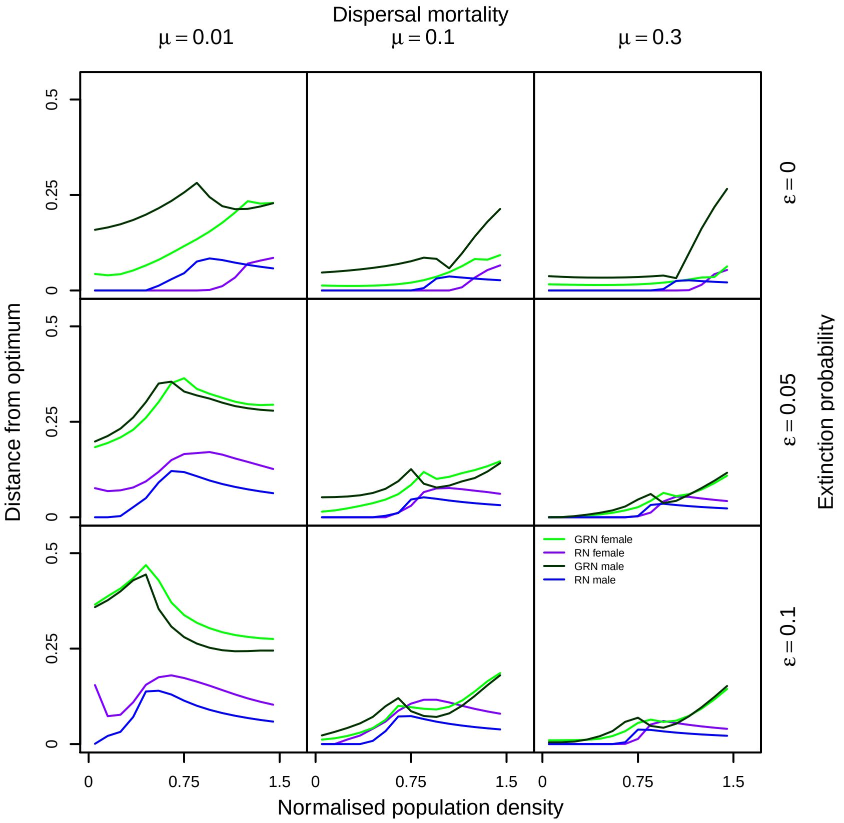


Figure S5: Average distance from optimal plastic response as function of normalised population density for GRN and RN models for DDD + sex bias. Dipersal mortality increases from left to right ($\mu \in \{0.01, 0.1, 0.3\}$), from top to bottom, extinction probability increases ($\epsilon \in \{0, 0.05, 0.1\}$). The optimal plastic response is calculated from the median of the evolved threshold $C_{thresh,male}$ and $C_{thresh,female}$ in the RN model. The plastic response for 1000 randomly chosen individuals in the GRN and RN models at end of the equilibrium metapopulation phase (20000 time steps) are evaluated at different normalised population densities $0, 0.1, ...1.5$, and the root mean squared distance is calulculated as a measure of deviation from this optimum. Similar to when on only DDD evolves, greatest distance from optimum in the GRN model is when dispersal mortality is low. Fixed parameters: $\lambda_0 = 2$ and $\alpha = 0.01$. Number of regulatory genes $n = 4$.

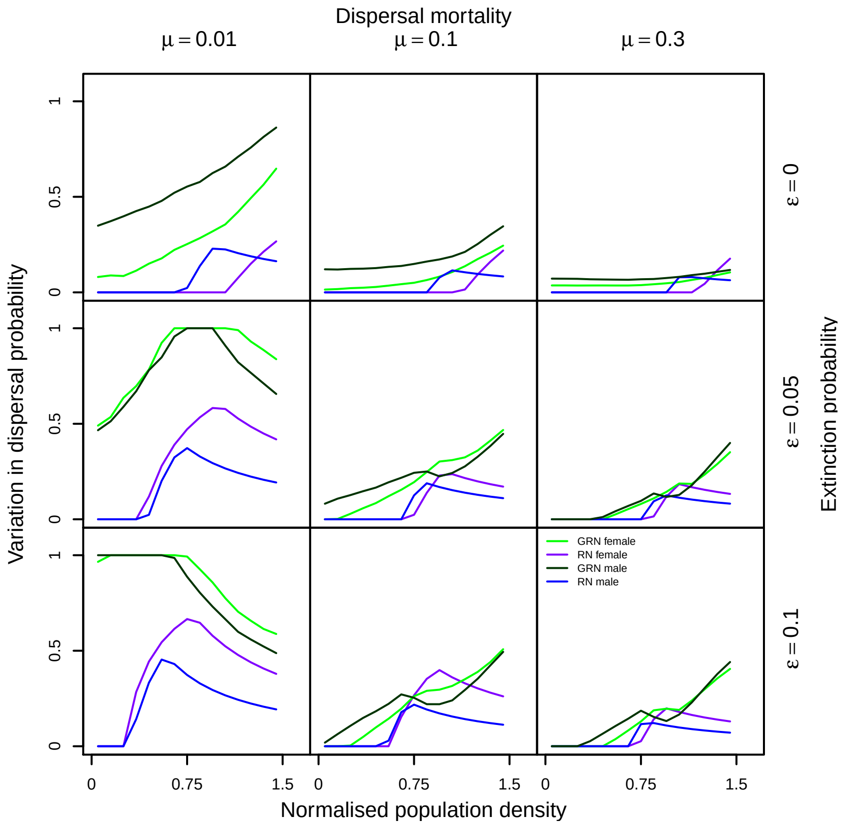


Figure S6: Phenotypic variation maintained in the GRN vs. RN model for DDD + sex bias as a function of normalised population density. Dipersal mortality increases from left to right ($\mu \in \{0.01, 0.1, 0.3\}$), from top to bottom, extinction probability increases ($\epsilon \in \{0, 0.05, 0.1\}$). Difference between the 95th and 5th percentile in dispersal phenotype as a function of normalised population density plotted for the GRN and RN model. Fixed parameters: $\lambda_0 = 2$ and $\alpha = 0.01$. Number of regulatory genes $n = 4$.

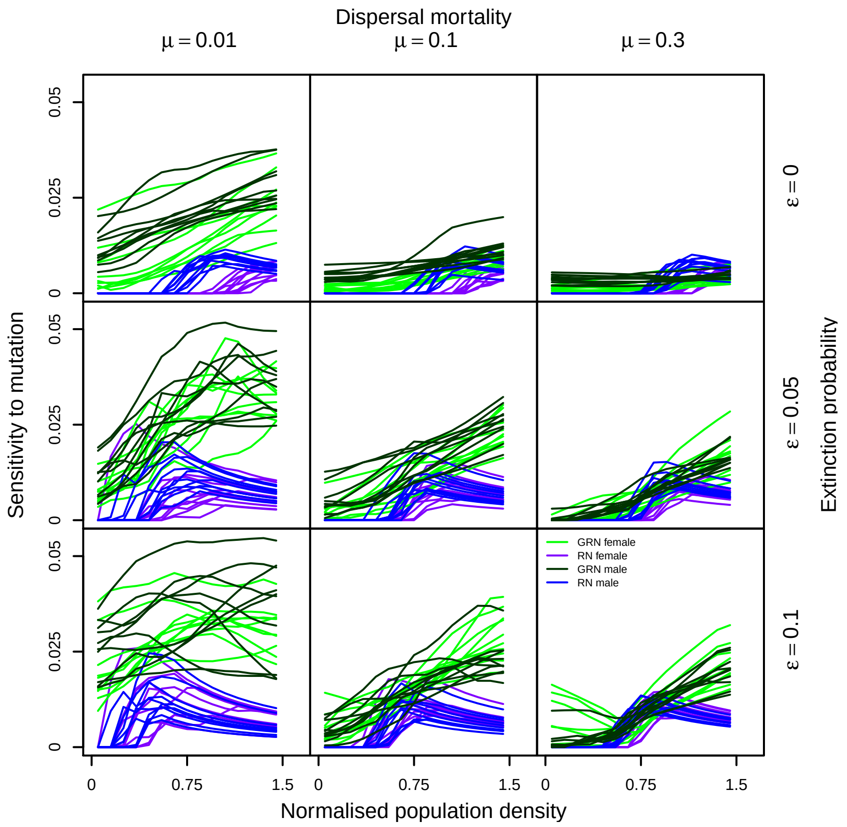


Figure S7: Sensitivity to mutation in the GRN and RN model for DDD + sex bias as a function of normalised population density. Dispersal mortality increases from left to right ($\mu \in \{0.01, 0.1, 0.3\}$), from top to bottom, extinction probability increases ($\epsilon \in \{0, 0.05, 0.1\}$). In both models, 1000 individual genotypes are sampled from the last time step of the equilibrium metapopulation phase ($t = 20000$). In the RN model, a perturbation drawn from a Gaussian distribution with mean 0 and standard deviation 0.1 is added to the evolved threshold $C_{thresh,male}$ and $C_{thresh,female}$ with probability 0.01. In the GRN model, a perturbation with the same mean and standard deviation is added to to an individual's locus with probability 0.01. This makes both models comparable, since per locus perturbation rate and effect are the same. The sensitivity to mutation is then calculated as the root mean squared difference between the phenotype evaluated from the perturbed and unperturbed genotype. Similar to the model for DDD alone, GRNs are more sensitive to mutation at low dispersal mortality. Fixed parameters: $\lambda_0 = 2$ and $\alpha = 0.01$. Number of regulatory genes $n = 4$.

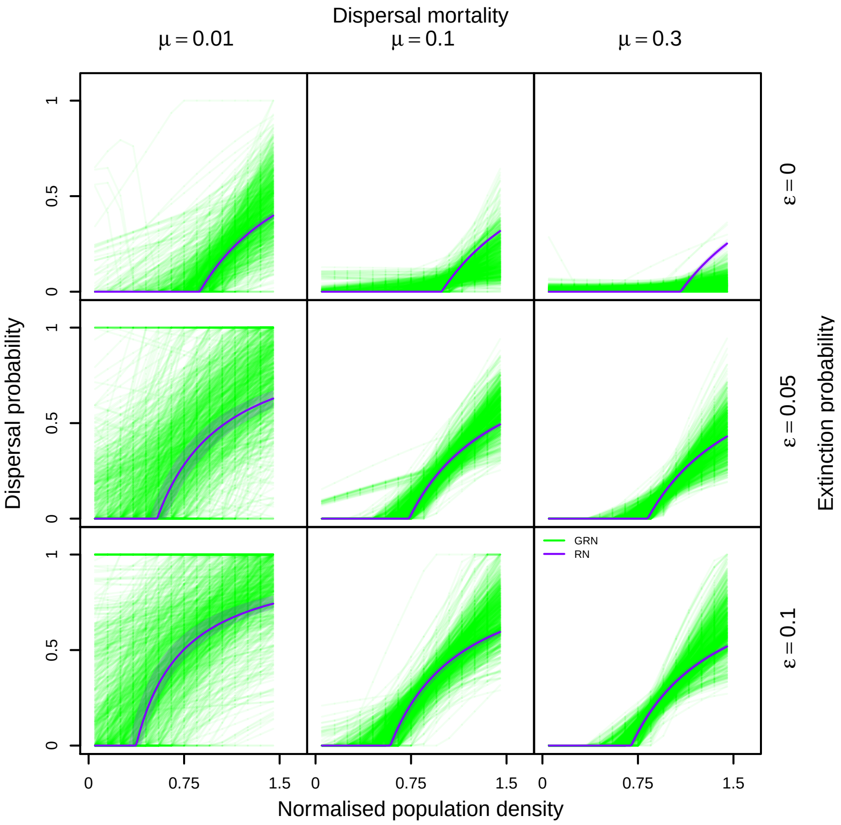


Figure S8: Density-dependent dispersal plastic responses in the GRN vs, RN model in the equilibrium metapopulation range core before the beginning of range expansion shown across the range of possible population densities. Dispersal mortality increases from left to right ($\mu \in \{0.01, 0.1, 0.3\}$), from top to bottom, extinction probability increases ($\epsilon \in \{0, 0.05, 0.1\}$). The green lines show the GRN density-dependent dispersal plastic response 1000 sampled GRNs pooled across 50 replicates in the range core before range expansions begin, whereas the purple lines show the expected plastic response from the RN model. We see that when we also depict plastic responses at population densities that do not occur frequently in equilibrium metapopulation conditions (unlike in Fig. 2 , where only those population densities are shown which occur frequently in equilibrium metapopulation conditions), there is a greater diversity of plastic responses maintained. Fixed parameters: $\lambda_0 = 2$ and $\alpha = 0.01$. Number of regulatory genes $n = 4$.

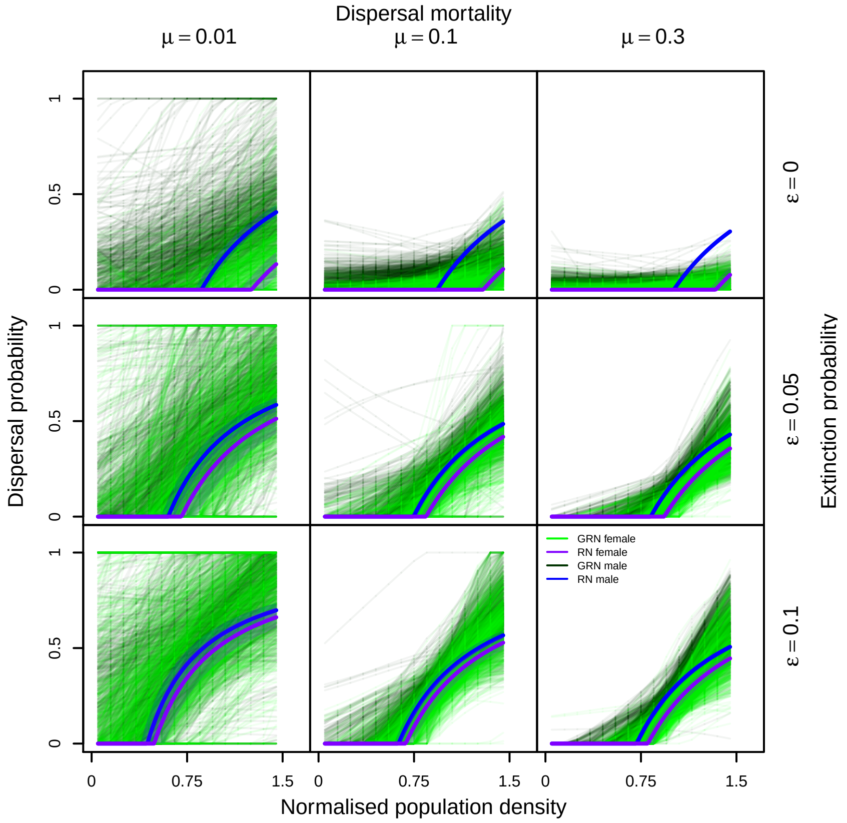


Figure S9: Density-dependent and sex-biased dispersal plastic responses in the GRN vs, RN model in the equilibrium metapopulation range core before the beginning of range expansion shown across the range of possible population densities. Dispersal mortality increases from left to right ($\mu \in \{0.01, 0.1, 0.3\}$), from top to bottom, extinction probability increases ($\epsilon \in \{0, 0.05, 0.1\}$). The dark green and green lines show the GRN density-dependent dispersal plastic response if male and female respectively for 1000 sampled GRNs pooled across 50 replicates, whereas the blue and purple lines show the expected plastic response from the RN model for males and females, respectively. We see that when we also depict plastic responses at population densities that do not occur frequently in equilibrium metapopulation conditions (unlike in main text Fig. 3, where only those population densities are shown which occur frequently in equilibrium metapopulation conditions), there is a greater diversity of plastic responses maintained. Fixed parameters: $\lambda_0 = 2$ and $\alpha = 0.01$. Number of regulatory genes $n = 4$.

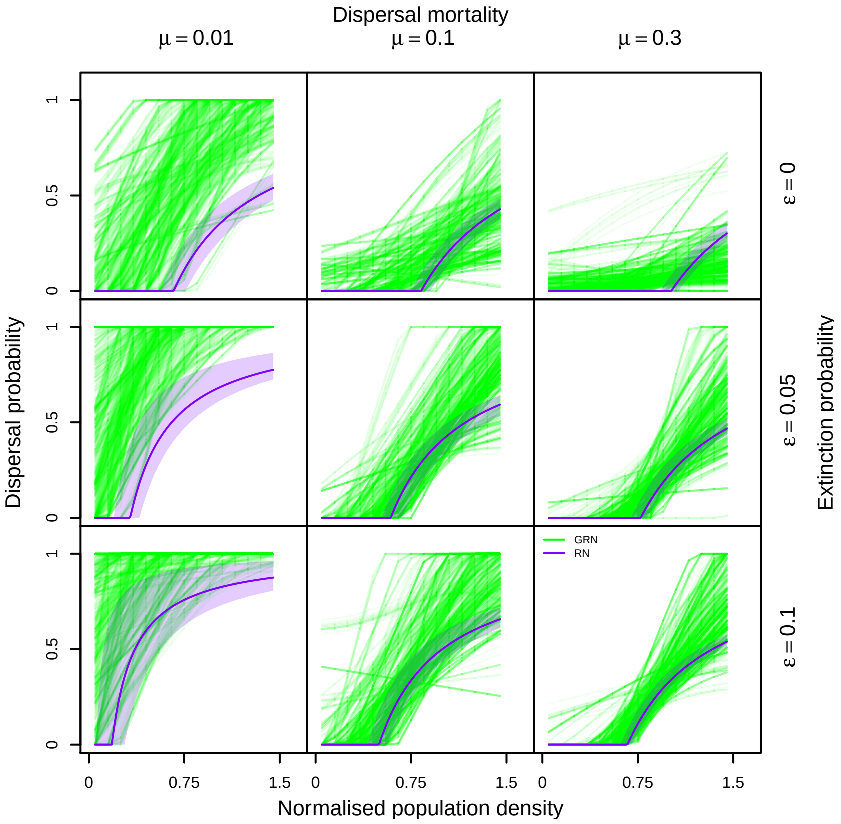


Figure S10: Density-dependent dispersal plastic responses in the GRN vs. RN model in the range front at the end of range expansion. Dispersal mortality increases from left to right ($\mu \in \{0.01, 0.1, 0.3\}$), from top to bottom, extinction probability increases ($\epsilon \in \{0, 0.05, 0.1\}$). The green lines show the GRN density-dependent dispersal plastic response for 1000 sampled GRNs pooled across 50 replicates, whereas the purple lines show the expected plastic response from the RN model. Fixed parameters: $\lambda_0 = 2$ and $\alpha = 0.01$. Number of regulatory genes $n = 4$.

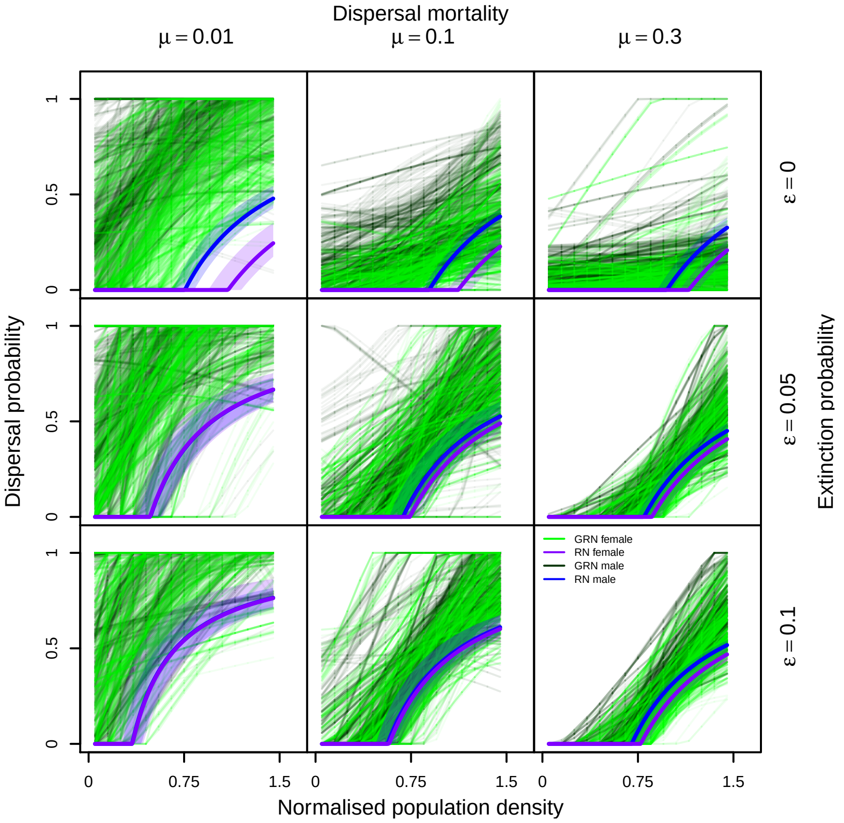


Figure S11: Density-dependent and sex-biased dispersal plastic responses in the GRN vs. RN model in the range front at the end of range expansion. Dispersal mortality increases from left to right ($\mu \in \{0.01, 0.1, 0.3\}$), from top to bottom, extinction probability increases ($\epsilon \in \{0, 0.05, 0.1\}$). The dark green and green lines show the GRN density-dependent dispersal plastic response if male and female, respectively for 1000 sampled GRNs pooled across 50 replicates, whereas the blue and purple lines show the expected plastic response from the RN model for males and females, respectively. Fixed parameters: $\lambda_0 = 2$ and $\alpha = 0.01$. Number of regulatory genes $n = 4$.

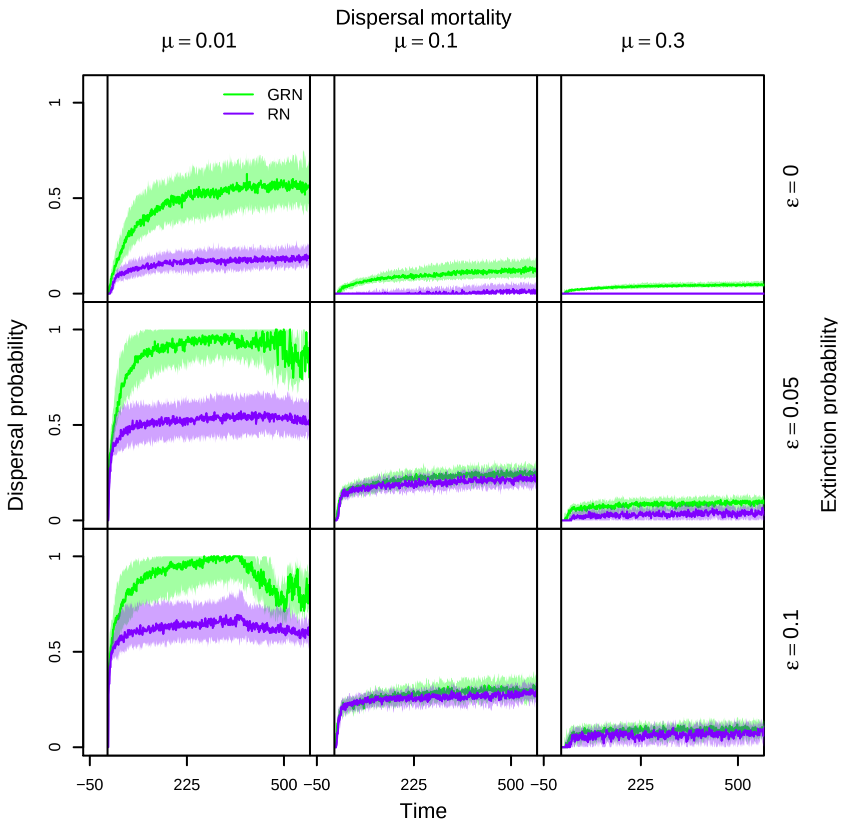


Figure S12: Change in dispersal phenotype in the GRN DDD vs, RN DDD model at the range front. Dispersal mortality increases from left to right ($\mu \in \{0.01, 0.1, 0.3\}$), from top to bottom, extinction probability increases ($\epsilon \in \{0, 0.05, 0.1\}$). Dispersal evolves to greater values in the GRN DDD model. Fixed parameters: $\lambda_0 = 2$ and $\alpha = 0.01$. Number of regulatory genes $n = 4$.

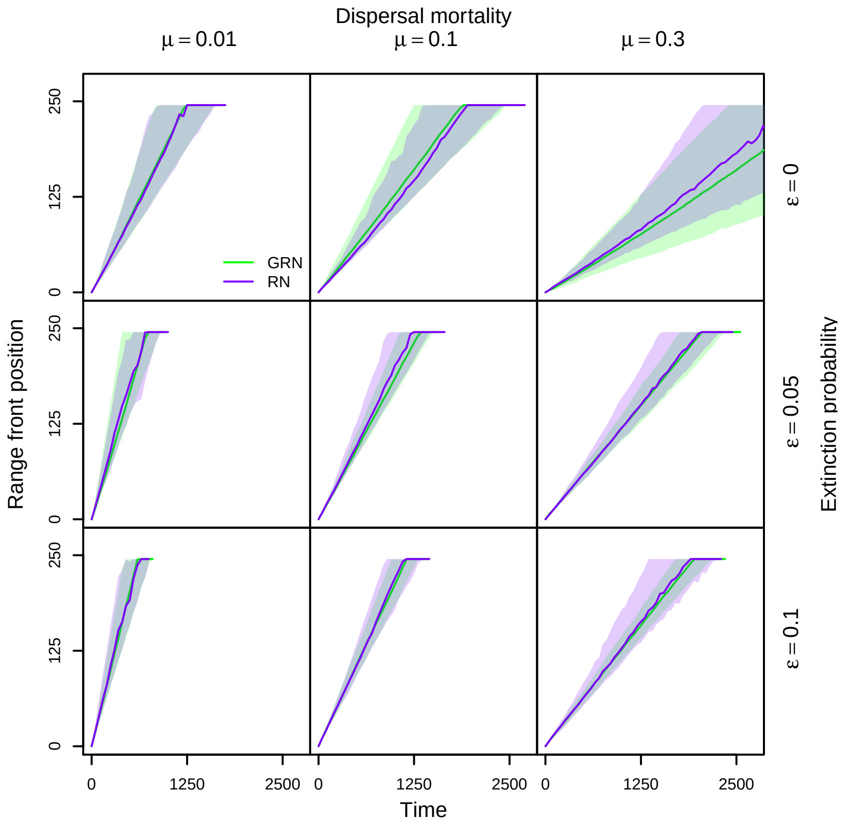


Figure S13: Sensitivity of range expansion dynamics to smaller mutation effects. Dispersal mortality increases from left to right ($\mu \in \{0.01, 0.1, 0.3\}$), from top to bottom, extinction probability increases ($\epsilon \in \{0, 0.05, 0.1\}$). We plot the range expansion dynamics for smaller per locus per allele mutation effects in the GRN DDD model ($(1/4)\sigma_m$). We find that there is no difference between range expansion dynamics in the two models. Fixed parameters: $\lambda_0 = 2$ and $\alpha = 0.01$. Number of regulatory genes $n = 4$.
